# Sensory experience and mTORC1 interplay orchestrates the maturation of cortical interneuron connectivity and tactile sensitivity

**DOI:** 10.64898/2026.07.22.740050

**Authors:** Clara A. Amegandjin, Maria Isabel Carreño-Muñoz, Ruggiero Francavilla, Jorelle Linda Damo Kamda, Marie Roussel, Antônia Samia Fernandes do Nascimento, Antoine Farley, Pegah Chehrazi, Alessandra Ciancone Chama, Bidisha Chattopadhyaya, Graziella Di Cristo

## Abstract

Sensory abnormalities are core features of neurodevelopmental disorders, including autism. Although interneuron dysfunction is hypothesized to contribute to these deficits, the underlying mechanisms remain unclear. Here, we demonstrate that mTORC1 dysregulation in parvalbumin-expressing (PV) interneurons drives heightened tactile exploration and defensiveness. These behavioral changes coincide with whisker-evoked cortical responses characterized by increased power but degraded temporal precision. Excitatory inputs to PV cells, their intrinsic excitability and *in vivo* firing rate during tactile exploration are reduced, suggesting that mutant PV cells are hypoactive. Whisker trimming restricted to the third postnatal week prevented mTORC1 hyperactivation, PV cell input and output connectivity deficits as well as abnormal tactile cortical responses and behavior in adult mutant mice. Further, this manipulation rescued sociability deficits. Altogether, these data suggest that the interplay between mTORC1 signaling and sensory experience in PV cells regulates their connectivity, and contributes to the proper development of tactile and social behavior.

## INTRODUCTION

The brain constantly perceives, processes and interprets sensory inputs, in turn determining appropriate behavioral outputs to successfully navigate the environment. Atypical sensory processing can interfere with a person’s ability to participate in everyday life, by affecting learning, coordination, language and behavior. For example, individuals struggling to process and filter out sensory stimuli appropriately may experience sensory overload or may be trigged easily by the environment, which might result in withdrawal, avoidance behavior and exacerbate stress and anxiety. On the other hand, individuals with low sensory registration may experience lack of awareness, lethargy and depressed mood. While the prevalence of sensory processing disorders ranges from 5-16% of children in the general population^1,2^, it is significantly higher in individuals with neurodevelopmental disorders (NDDs), such as autism spectrum disorders^3,4^ (ASD). Atypical sensory processing was indeed recently included as one of the core features of ASD in the DSM-5, since it is estimated that up to 90% of ASD individuals have atypical sensory experiences^3^. In particular, recent studies suggest that abnormal responses to touch are highly correlated with, and predictive of, ASD severity^5,6^. During development, sensory experience plays a critical role in shaping brain circuits, by regulating synaptic consolidation and pruning during sensitive postnatal periods. It is therefore conceivable that atypical processing of sensory experience during early development may alter the developmental trajectory of neural circuits, thereby contributing to the cognitive and social behavioral alterations associated with ASD and related NDDs^7^. How impaired circuit connectivity and function contribute to atypical sensory processing —and whether postnatal sensory modulation can prevent these deficits —remains to be determined.

GABAergic neurotransmission plays a key role in sensory function, including tactile stimuli processing^8^. Further, GABAergic circuit alterations have been observed in both animal models of and individuals with ASDs^9^, leading to the hypothesis that differences in inhibition/excitation within cortical circuits contributes to their symptomatology^8–10^. Among GABAergic neuron subtypes, parvalbumin (PV)-expressing basket interneurons (hereafter, PV cells) preferentially form synapses onto the soma and proximal dendrites of their postsynaptic partners. By providing fast perisomatic inhibition, they exert precise control over pyramidal neuron output and network oscillations. Dysfunction of PV cells is strongly associated with ASD, both in human and across genetically distinct mouse models of NDDs characterized by high ASD prevalence^9^. In particular, PV circuit hypofunction has been causally linked to abnormal processing in different sensory domains in NDD mouse models^11–13^. Consequently, much effort has been recently devoted to explore the cellular mechanisms leading to PV cells hypofunction in NDDs with ASD.

The serine/threonine protein kinase mechanistic target of rapamycin complex 1 (mTORC1) is a signalling hub, integrating environmental information regarding energy availability and stimulating anabolic molecular processes and cell growth^14^. Consistent with its multifaceted roles in the nervous system, deregulation of mTORC1 signalling is associated with numerous NDDs^14,15^. In particular, mutations in genes encoding mTORC1 regulators lead to NDDs with shared neurological manifestations, collectively referred as “mTORopathies”, which are characterized by the high prevalence of ASD^14,16^. Recent studies suggest that cortical PV cell maturation is impaired by excessive mTORC1 signaling^17,18^, nevertheless, whether mTORC1 dysregulation specifically in PV cells contributes to sensory processing abnormalities is not known. Furthermore, since PV cell maturation is significantly shaped by sensory experience^19–21^, it is conceivable that sensory experience interacts with mTORC1 dysregulation to determine PV cell impairment, and possibly sensory processing dysfunction, in adulthood.

Here, we tested this hypothesis using a mouse model of *Tsc1*-haploinsufficiency restricted to PV cells. Tsc1, together with Tsc2 and Tbc1d7, forms a complex that negatively regulates mTORC1 signaling. PV cell-specific conditional *Tsc1* heterozygous mice show behavioral sensory phenotypes, including increased voluntary exploration of textured objects, suggestive of a tactile seeking behavior, and increased tactile defensiveness behavior following passive whisker stimulation, suggesting tactile hyper-reactivity. Mutant mice further show abnormal cortical activity in the primary somatosensory cortex during free object exploration and following passive whisker stimulation. These phenotypes are rescued by treatment with the mTORC1 inhibitor, rapamycin, during the third postnatal week, suggesting that mTORC1 hyperactivity in PV cells during this sensitive developmental window has a long-lasting impact on sensory processing. At the cellular level, mutant PV cells show reduced cortical glutamatergic inputs in adult primary somatosensory cortex, as indicated by immunolabeling, electron microscopy and *ex vivo* patch-clamp recording. Further, the firing rate of narrow-spiking neurons (putative PV cells) is reduced in freely exploring adult mice, suggesting that *Tsc1* haploinsufficient PV cells are hypoactive. PV cell input and output connectivity, state-dependent spontaneous and whisker-evoked cortical activity, and tactile behaviors are rescued by whisker trimming during the third postnatal week in mutant adult mice. Taken together, these data suggest that the interaction between mTORC1 signaling dysregulation and sensory experience during a critical time window of PV cell maturation leads to cortical activity and tactile behavioral abnormalities in adulthood. Finally, whisker deprivation restricted to the third postnatal week prevents the development of sociability impairments in adult mutant mice, indicating a causal link between impairments in early tactile processing and social dysfunction in mice.

## MATERIALS AND METHODS

### Animals

*Tsc1* floxed mice with loxP sites flanking exons 17 & 18 of *Tsc1* gene (*Tsc1^flox/flox^*) were purchased from Jackson Laboratories (Cat# 005680). Two driver mouse lines expressing Cre recombinase, *PV-Cre* (Jackson Laboratory, RRID:IMRS_JAX:017320) and *Nkx2.1-Cre* (Jackson Laboratories, RRID:IMRS_JAX:008661) were crossed to the *Tsc1* floxed mice and the respective progenies were backcrossed to generate the heterozygous and control genotypes within the same litter. To control for the pattern of expression of Cre, we introduced the RCE allele using Gt(ROSA)26Sortm1.1(CAG-EGFP)Fsh/J mice (Jackson Laboratories; RRID:MMRRC_032037-JAX). The RCE line carries a loxP-flanked STOP cassette upstream of eGFP sequence within the Rosa26 locus. Removal of the loxP-flanked STOP cassette by Cre-mediated recombination allows promoter-specific downstream eGFP expression^74^. All mice were housed under standard pathogen-free conditions in a 12h light/dark cycle with *ad libitum* access to sterilized laboratory chow diet. Animals were treated in accordance with the Canadian Council on Animal Care and protocols were approved by the Animal Care Committee of CHU Ste-Justine Research Center.

### Mice Genotyping

DNA was extracted from mouse tail and genotyped to detect the presence of Cre alleles and Tsc1 conditional and wild-type alleles. Polymerase chain reaction (PCR) was performed using 3 separate primers: F4536 (5’-AGGAGGCCTCTTCTGCTACC-3’), R4830 (5’-CAGCTCCGACCATGA AGTG -3’) and R6548 (5’-TGGGTCCTGACCTATCTCCTA-3’) with band sizes of 295bp for the wild-type and 480bp for the floxed allele. Three separate primers were also used for detecting Cre in *PV*-Cre mice: F1 (5’-CAGCCTCTGTTCCACATACACTCC-3’), F2 (5’- GCTCAGAGCCTCCATTCCCT-3’) and R1 (5’- TCACTCGAGAGTACCAAGCAGGCAGGA GATATC-3’), which generated 400bp and 526bp (mutant and wild-type) bands. To detect the presence of RCE alleles, 3 separate primers were used: RCE-Rosa1(5’-CCCAAAGTCGCTCTGAGTTGTTATC-3’), RCE-Rosa2 (5’GAAGGAGCGGGAGAAATGGATATG-3’) and RCE-Cag3(5’-CCAGGCGGGC CATTTACCGTAAG-3’), which generated 350bp and 550bp bands. Finally, three separate primers were used for detecting Cre in the Tg(Nkx2.1-Cre) breeding, which generated 550 and 220 bp (mutant and wild-type) bands^17^.

### Mouse behavior tests

Behavioral tests were conducted during the light-on period of the day, utilizing age-matched (P60-P80) littermates. To habituate the mice to the experimenter, mice were handled at least 2 min per day, 5 times per week, for the two weeks prior to testing. Investigators were blind to genotype during both testing and analysis.

#### Texture discrimination test

The test was performed in dimmed light the open-field apparatus, with a camera mounted above the arena, using age-matched male littermates. This test consisted of three trials, habituation, learning and testing phases (Supplemental Figure 1A). To mitigate olfactory cues, all objects (familiar and new) and the box were cleaned with 70% ethanol between sessions. Between different animals the arena and all objects were cleaned with acetic acid (0.1%), and then with ethanol (70%). 3D-printed objects were identical in size and color, but different in texture (Supplemental Figure 1B). Mice were first habituated to the open field chamber by allowing free exploration of an empty chamber for 5 min (habituation); then removed from the testing arena and placed in a transport cage. After 5 minutes, the mouse was placed in the testing arena, equidistant to the two identical objects placed in the center of the arena. The two objects were positioned equidistant from the center of the arena, and equidistant from the walls of the arena. Each mouse was allowed to explore the objects for 10 minutes (learning phase). Animals were then removed from the testing arena and placed in a transport cage for 5 minutes. The mouse was then placed back into the chamber and allowed to explore two objects, one featuring the previously encountered texture and the other featuring a novel texture, for 10 minutes (testing phase)^22,23^. Investigation was defined as the act of positioning the nose within a distance of less than 2 cm from the object or making direct contact between the nose and the object. Activity such as resting, grooming in close proximity to the object were not considered instances of investigation. The amount of time the mouse spent investigating each of the objects was manually determined for all the trials and was used to calculate the discrimination index as well as the total exploration time. Total exploration time was calculated as (time spent investigating the novel object + time spent investigating the familiar object). The discrimination index was calculated as (time spent investigating the novel object-time spent investigating the familiar object)/ total exploration time. Mice were included in the analysis only if 1) they explored both objects in the learning and testing phases, and 2) if the total exploration time was > 10 sec.

#### Novel object recognition test

The novel object recognition was conducted following the same protocol described above, only the testing phase differed since the novel object introduced in that phase differed from the familiar object in shape and color, but the texture of both objects was identical.

#### Social behavior test

Between each mouse, the arena and the cages were cleaned with acetic acid (0.1%), and then with ethanol (70%). Mice of both sexes were placed in the middle of the central chamber and allowed to explore all the chambers for 10 min (habituation). Following cleaning with ethanol (70%), a wire cage containing an unfamiliar conspecific of the same sex and age (Stranger) was placed inside one chamber, while an empty wire cage was placed in the second chamber. Stranger mice originated from different home cages and had never been in physical contact with the test mice. Mice were allowed to freely explore the three chambers of the apparatus for 10 min. Social approach was evaluated by manually quantifying the time spent by the test mice with the object or the mouse in each chamber during the 10 min session. Investigation was defined as the act of positioning the nose within a distance of less than 2 cm from the empty cage or the stranger mouse or making direct contact between the nose and the cage/mouse. Activity such as resting, grooming in close proximity to of the object were not considered instances of investigation. Total exploration time was calculated as (time spent investigating the empty cage + time spent investigating the mouse). The discrimination index was calculated as (time spent investigating the mouse-time spent investigating the object)/ total exploration time).

#### Brush test

Mice were tested as previously described^26^. Briefly, mice were habituated in a small Plexiglas chamber on a wire mesh table for 1 hour the day prior to testing and for an additional 1h immediately before testing. The testing was performed using a paintbrush that was prepared by trimming its tip and by removing the outer layer of hairs. The brush was applied with minimal pressure to the planter surface of the testing paw. Testing alternated between the ipsilateral and contralateral paws with a 5 min gap between each application. The application was repeated 10 times and the average nocifensive response score was determined for each animal. Non-aversive behavior was defined by the absence of response or withdrawal of the paw without escape (score 0) and withdrawal of the paw with escape (score 1). Aversive behavior was defined by the presence of flipping, guarding of paw, or startle-like jumping (score 2), and multiple flinching responses or licking of the paw (score 3). For this test, the average score is close to zero for naïve mice, but above 1.5 (more often around 2.0) in mice with mechanical hypersensitivity (Cheng et al. 2017).

#### Tactile seeking exploration assay

This assay was performed in chronically implanted mice to analyze cortical activity within the primary somatosensory circuit during free whisking-based exploratory behavior. Starting 1 week after electrode implant (see *In vivo* electrophysiology section below), animals were habituated daily for 3 days to an open field (15–20 min each time). We used a dimly illuminated open field environment (45 by 35 cm), surrounded by 60 cm-high opaque walls and equipped with video monitoring on top of the arena. The open field’s floor and walls were sprayed and wiped clean with 70% ethanol 30 min before the introduction of each animal. On day 4, two identical textured objects (rougher textured object in Supplemental Figure 1B) were placed in the open field and both local field potential (LFP) and video were recorded for 30 minutes. Object exploration periods were automatically detected by using DeepLabCut algorithms. To build the DeepLabCut deep-learning network we manually labelled a variety of contact types: nose-object, body-object, rearing (two frontal paws on the object) and climbing (four paws on the object). Exploration bouts were defined as nose-object direct contact that lasted at least 0.2 s. We selected frames in which DeepLabCut detection showed a likelyhood >0.9. We also inspected visually about half of the videos to confirm detection accuracy. In addition, Deeplabcut analysis provided continuous XY coordinates that enabled quantification of locomotor activity by calculating Euclidean distances of the body center over frames. Locomotor analysis included total time moving (sum of all running periods, in seconds, when body center moved > 1.5 cm/s for at least 0.5 s), total distance moved (cm), mean velocity (cm/s) and the identification of immobility periods (when body center moved <1.5 cm/s for at least 1 s).

#### Tactile defensiveness assay in head-restrained mice

Tactile defensiveness was assessed for each mouse during whisker stimulation sessions, as described in He et al^41^. Mice were habituated for 7–10 days to the head-fixed system situated on an air-supported ball treadmill. The habituation period was progressively longer, starting from 5 up to 25 min daily. Once mice were habituated to staying on the ball for 25 min, they were recorded during 3 minutes of baseline activity and three different whisker stimulation protocols (see *In vivo* electrophysiology section below). The total duration of each session was 20-25 min. For every session, we analyzed the videos and manually scored the durations of defensive behaviors (grabbing the stimulator) and adaptive behaviors (grooming). The total time spent on each behavior, as well as the total session time, was summed across all sessions for each mouse. Behavioral percentages were then calculated by dividing the total time spent on each behavior by the total session time.

### Rapamycin treatment

Rapamycin (3 mg/kg; i.p.) was administered daily to pups from P14 to P21. Rapamycin stock solution (20 mg/ml in 100% ethanol) was stored at −20°C. Before injection, stock solution was diluted in 5% Tween 80 and 5% polyethylene glycol 400 to a final concentration of 1 mg/ml rapamycin in 4% ethanol (Buckmaster and Wen 2011). Control mice were injected with the vehicle solution.

### Sensory deprivation

The transient restriction of neonatal vibrissal input was achieved by trimming the whiskers close to the intersection with the skin of P14 pups. Mice were restrained by hand and all whiskers were trimmed every two days to within 1 mm of the skin for 8 days (P14 to P21). Pups were then allowed to mature without further intervention except for weekly cage cleaning. Before behavioral test, all whiskers regrew and were of the equivalent length to untrimmed mice. For social behavioral testing (Figure 7), sham mice were restrained by hand and the scissor was manoeuvred close to the whiskers without touching them every two days from P14 to P21.

### Immunohistochemistry

Mice of both sexes were perfused transcardially with saline followed by 4% Paraformaldehyde (PFA 4%) in phosphate buffer (PB 0.1M, pH 7.2). Brains were post-fixed with 4% PFA overnight and subsequently transferred to a 30% sucrose solution in sodium phosphate-buffer (PBS) for 48hrs. They were then frozen in molds filled with Tissue Tek using a 2-Methylpentane bath cooled with a mixture of dry ice and ethanol (∼ −70°C). Optimal cutting temperature and coronal sections of 40 µm were obtained using a cryostat (Leica VT100). Brain sections were blocked in 10% normal goat serum (NGS) and 1% Triton X-100 for 2 hr at RT. Slices were then incubated for 48h at 4°C with the following primary antibodies: mouse anti-PSD-95 (1:500, Invitrogen, Cat# MA1045), rabbit anti-Vglut1 (1:100, Invitrogen, Cat# 482400), mouse anti-PV (1:1000, Swant, Cat# 235), rabbit anti-PV (1:5000, Swant, Cat# PV27), mouse anti-gephyrin (1:500, Synaptic Systems, Cat# 147021), chicken anti-GFP (1:1000, Abcam, Cat# 13970); rabbit anti-phospoS6 (1:1000, Cell Signaling, Cat# 5364); chicken anti-SynCAM1(1:500, MBL Life Science, Cat# CM004-3); rabbit anti-Nlgn3 (1:1000, Synaptic Systems, Cat# 129113). It was followed by incubation with secondary antibodies for 2h at RT to visualize primary antibodies. The secondary antibodies used were Alexa-Fluor conjugated 488, 555, 633, and 647 (1:400, Life technologies; 1:1000, Cell Signaling Technology). After rinsing in PBS (three times), the slices were mounted in Vectashield mounting medium (Vector).

### Confocal Imaging and Quantitative analysis

All imaging was performed using Leica confocal microscopes (SP8 or SP8-STED). We imaged layer 5 of the primary somatosensory cortex (barrel cortex) using 63X glycerol-immersion objective (NA1.3), Zoom 1.5. At least three confocal stacks from 3 different brain sections were acquired in layer 5 for PV/gephyrin, Vglut1/PSD-95 and Nlgn3/PSD95 puncta, and for SynCAM1/PV labeling, with z-step sizes of 0.5 µm. Vglut1+, PSD-95+, Vglut1+/PSD-95+, and Nlgn3/PSD95 puncta were counted around GFP positive somata after selecting the confocal plane with the highest soma circumference using ImageJ-Fiji software (custom-made ImageJ-Fiji macro). PV+, gephyrin+, PV+/gephyrin+, and Nlgn3/PSD95 pucta were counted around putative pyramidal cell somata as well. To quantify the number of puncta, images were exported as TIFF files and analyzed using Fiji (Image J) software. We first manually outlined the profile of each cell soma and used a series of custom-made macros in Fiji as previously described (Chehrazi, Lee et al. 2023). Briefly, after applying subtract background (rolling value = 10) and Gaussian blur (σ value = 2) filters, the stacks were binarized and the colocalized puncta were independently identified around the perimeter of a targeted cell after selecting the focal plane with the highest soma circumference. At least 4 innervated somata were selected in each confocal image. Puncta were quantified after filtering particles for size (included between 0 and 2 μm2) and circularity (included between 0 and 1). Puncta density for each mouse was normalized over the mean of the control group for each experiment.

Fluorescence intensity of pS6 signal in layer 5 PV cells was calculated using ImageJ software. Eight to ten PV+ cells were chosen in each confocal stack and their cytoplasm was encircled by using the polygon tool. We then calculated the mean gray value of pS6 signal of each selected PV cells and subtracted the background. Background mean signal was calculated in at least three regions devoid of any staining from the same focal plane where the selected PV cells were located.

Quantification of SynCAM1 expression levels in dendrites and neuropile was performed as previously described (Ribic, Crair et al. 2019). Briefly, integrated density of SynCAM1 was measured around PV cells dendrites and in the neuropile of somatosensory cortex (Layer 5) using the quantification tools in Leica LasX software. The values were then normalized by subtracting the average intensities of the background. 6 to 8 dendritic segments were quantified on average from each animal from 3-4 brain sections. All quantification were done by investigators blind to the genotype or experimental conditions.

### Electron microscopy

The electron microscopy was carried out on 2 groups, *Tsc1*^Ctrl^ and *PV*-*Cre*;*Tsc1^flox/+^* mice, at P60. Mice were anesthetized and perfused with 0.1M PBS (0.9% NaCl in PB 0.2M, pH 7.4) followed by 2,5% glutaraldehyde + 2% PFA in 0.1M PB, pH 7.4. Following perfusion, the brains were further fixed for 2 hours at room temperature (RT) in the perfusion solution. Transverse 50-µm-thick sections of the brain were cut in cooled PBS with a vibratome (Leica, VT1000S). They were stored at −20°C in antifreeze solution (40% PB, 30% ethylene glycol, 30% glycerol) until used. Sections were immersed in 0.1% borohydride (in PBS) for 15 min at room temperature (RT), washed in PBS, and processed freely floating following a pre-embedding immunoperoxidase protocol previously described (Tremblay et al. 2007). Briefly, after rinsing in PBS, sections were preincubated (1-hour) at RT in a protein blocking solution (Expose Rabbit-Specific HRP/DAB detection IHC Kit, Abcam, Cambridge, UK, ab80437). Then, the sections were incubated for 48 hours at 4°C with rabbit anti-PV (1:1000, Swant, Cat# PV27) in PBS containing 1% NGS, followed by wash (three times in PBS) and incubation for 45 min at RT, in goat anti-rabbit horseradish peroxidase (HRP) conjugate (Abcam, Expose Kit, Cat# ab80437). After rinsing in PBS, immunoreactivity was visualized with hydrogen peroxide in the presence of diaminobenzodine (DAB Chromogen, Abcam Expose Kit, Cat# ab80437). Thereafter, sections were rinsed in PBS, postfixed flat in 1% osmium tetroxide for 1 hour and dehydrated in ascending concentrations of ethanol (50%, 70%, 90%, 100%, and finally in ethanol anhydrous). They were then treated with propylene oxide and then impregnated in resin overnight (Durcupan ACM; Sigma) at RT, mounted on aclar embedding film (EMS, Hatfield, PA) and cured at 55°C for 48 hours. Areas of interest from the somatosensory cortex (layers 5/6, barrel cortex) were excised from the embedded sections and glued to the tip of prepolymerized resin blocks. Ultrathin (50-70 nm) sections were cut with an ultramicrotome (Reichart UltracutS, Leica, Wetzlar, Germany), collected on bare 150 square-mesh copper grids (Electron Microscopy Sciences, Hatfield, PA), stained with lead citrate, and examined at 80 KV with a Philips CM100 electron microscope, equipped with an 8 MB digital camera (AMT XR80).

To analyze the electron microscopy data, cellular profiles were identified according to well established criteria^75^. All PV labeled structures were classified in different categories such as: dendritic shafts, axons and axon terminals. All the subcellular profiles that were difficult to identify were classified as “unknown”. To provide a better appraisal of the frequency of each type of cellular elements displaying immunolabelling, about eighty to hundred micrographs were randomly taken at 25000X in each animal, corresponding to a total surface of ∼ 2000 µm^2^. Labeled profiles were counted in all micrographs. Results were expressed as number of immunopositive profiles per 100 µm^2^ of neuropil then normalized over results from the control mice. The area of neuropil and synapses lengths were measured using Neurolucida (MicroBrighField).

### Ex Vivo Electrophysiology

Recording and data analysis were performed by investigators blind to the genotype or experimental conditions.

#### Slice Preparation and Patch-Clamp Recordings

Coronal slices (thickness, 350 μm) were prepared from somatosensory cortex of P60-85 *PVcre-Cre;RCE^f/f^;Tsc1^+/+^*(*Tsc1*^Ctrl^) and *PV-Cre;RCE^f/f^;Tsc1^f/+^*littermate males. Briefly, animals were anesthetized deeply with ketamine-xylazine mixture (ketamine: 100 mg/kg, xylazine: 10 mg/kg), transcardially perfused with 25 mL of ice-cold cutting solution (containing the following in mM: 250 sucrose, 2 KCl, 1.25 NaH_2_PO_4_, 26 NaHCO_3_, 7 MgSO_4_, 0.5 CaCl_2_, and 10 glucose, pH 7.4, 330–340 mOsm/L) and decapitated. Slices were cut in the cutting solution using a vibratome (VT1000S; Leica Microsystems), and transferred to a heated (37.5°C) oxygenated recovery solution containing the following (in mM): 124 NaCl, 2.5 KCl, 1.25 NaH_2_PO_4_, 26 NaHCO_3_, 3 MgSO_4_, 1 CaCl_2_, and 10 glucose; pH 7.4; 300 mOsm/L, and allowed to recover for 45 min. During experiments, slices were continuously perfused (2 mL/min) with standard artificial cerebrospinal fluid (ACSF) containing the following (in mM): 124 NaCl, 2.5 KCl, 1.25 NaH_2_PO_4_, 26 NaHCO_3_, 2 MgSO_4_, 2 CaCl_2_, and 10 glucose, pH 7.4 saturated with 95% O2 and 5% CO2 at near physiological temperature (30–33°C). PV-positive interneurons (PV+) located in layer 5 of barrel cortex were visually identified as eGFP-expressing cells under an epifluorescence microscope with blue light (filter set: 450–490 nm). All electrophysiological recordings were carried out using a 40x water-immersion objective. Recording pipettes were pulled from borosilicate glass (World Precision Instruments) with a PP-83 two-stage puller (Narishige) to a resistance range of 5–7 MΩ when backfilled with intracellular solution. Whole-cell patch-clamp recordings from PV+ interneurons were performed in voltage-clamp mode. For this recording we used an intracellular Cs+-based solution containing (in mM): 130 CsMeSO_4_, 5 CsCl, 2 MgCl_2_, 10 phosphocreatine, 10 HEPES, 0.5 EGTA, 4 ATP-TRIS, 0.4 GTP-TRIS, 0.3% biocytin, 2 QX-314 (pH 7.2–7.3; 280–290 mOsm/L). Voltage-clamp recordings were performed to analyze the excitatory drive received by PV + cells. Series resistance in voltage-clamp were monitored throughout the experiment and cells with series resistance changes >15% were excluded. Recordings of miniature excitatory postsynaptic currents (mEPSCs) were performed in voltage-clamp at –70 mV in the presence of tetrodotoxin (TTX; 1 μM; Alomone Labs). For current-clamp recordings we used an intracellular K^+^-based solution containing (in mM): 130 KMeSO4, 2 MgCl2, 10 di-Na-phosphocreatine, 10 HEPES, 4 ATP-Tris, 0.4 GTP-Tris, and 0.3% biocytin (Sigma), pH 7.2–7.3, 280–290 mOsm/L. Passive and active membrane properties were analyzed in current clamp mode. Active membrane properties and frequency-current (F-I) relationship for evoked firing were recorded by subjecting cells to multiple current step injections (step size 40 pA) of varying amplitudes (–200 to 600 pA). Passive membrane properties (resting membrane potential, input resistance, and membrane capacitance) were obtained immediately after membrane rupture. Input resistance was calculated as the slope of the linear fit to subthreshold steady state membrane voltage values and the injected current (V = a + b*I, where V is the steady state membrane voltage, I is the injected current, and b is the input resistance). Membrane potentials were maintained at –80 mV, series resistances (10–18 MΩ) and input resistances were monitored on-line with a 40pA current injection (150 ms) given before each 500 ms current injection stimulus. Only cells with resting membrane potential more negative than −60mV at the start of recording and spikes with overshoot were considered for further analysis. We performed bridge balance and neutralized the capacitance before starting every recording. The bridge balance was monitored throughout the experiment, and neurons showing changes of >15% in bridge balance during the recording were discarded. Data acquisition (filtered at 2–3 kHz and digitized at 10 kHz; Digidata 1440, Molecular Devices, CA, United States) was performed using the Multiclamp 700B amplifier and the Clampex 10.6 software (Molecular Devices).

#### Data Analysis

Analysis of electrophysiological recordings was performed using Clampfit 10.7 (Molecular Devices). For the analysis of mEPSCs a minimum of 200 events were sampled per cell over a 2 min period using an automated template search algorithm in Clampfit. All events were counted for amplitude and frequency analysis. Charge transfer was calculated by integrating the area under the mEPSC waveform. For the analysis of current-clamp recordings from PV cells, rheobase was measured as the minimal current necessary to evoke an action potential (AP). For the analysis of AP properties, the first AP appearing within a 50ms time window from beginning of current pulse was analyzed. AP latency was measured as the time between current step onset and when membrane voltage reached AP threshold. The AP amplitude was measured from the AP threshold to the peak. The AP half-width was measured at the voltage level of the half of AP amplitude. The AP rise time was detected between the AP threshold and the maximal AP amplitude, while the AP fall time between the maximal AP amplitude and the AP end. The fast afterhyperpolarization (fAHP) amplitude was determined as the minimum voltage following the action potential peak subtracted from the action potential threshold. fAHP time was determined as the time between action potential threshold and the negative peak of fAHP. The hyperpolarization-activated cation current (Ih)-associated voltage rectification (Ih sag) was determined as the amplitude of the membrane potential sag from the peak hyperpolarized level to the level at the end of the hyperpolarizing step when the cell was hyperpolarized to –100 mV. The membrane time constant (τ) was measured offline using an exponential fit of voltage responses to negative hyperpolarizing step currents of –40 pA.

#### Anatomical identification and immunohistochemistry

For post-hoc anatomical identification, neurons were filled with biocytin (Sigma) during whole-cell recordings. Slices with recorded cells were fixed overnight with 4% PFA at 4 °C. To reveal biocytin, the slices were permeabilized with 0.3% triton X-100 and incubated at 4 °C with a streptavidin-conjugated Alexa-488 (1:1000) in TBS. Confocal images of biocytin-filled cells were obtained using a Leica TCS SP8 DLS imaging system coupled to a 488-nm Argon laser. Z-stacks were acquired with a 1-µm step.

### *In Vivo* Electrophysiology

#### Surgery

Mice were implanted between P70-P80. Chronic implants were performed under deep anaesthesia, induced by an intraperitoneal injection of Ketamine/Xylazine/Acepromazine cocktail (80 mg/kg Ketamine, 15 mg/kg Xylazine, and 2 mg/kg Acepromazine). A craniotomy was performed on the right side of the somatosensory cortex using the coordinates −1.5 mm posterior to Bregma, 2.7 mm lateral to the midline. Custom-made clusters of four to six 50 μm-diameter-insulated-tungsten wires were implanted. Wire tips (recording sites) were 50–150 μm apart, and the deepest was positioned 0.8 mm deep. A ground-and-reference wire was also gently introduced in the contralateral frontal lobe and a 2cm carbon-fibre bar was placed on the back of the skull. The whole structure was then daubed with dental acrylic to encase the electrode-microdrive assembly and anchor it to the skull. During the whole surgery, body temperature was maintained at 36°C–37°C by a thermostatically controlled heating pad. For 1 week after surgery, animals were treated with Metacam and the skin around the implant was disinfected daily for 3 days following the surgery. Electrode placement was confirmed at the end of each experiment by histological analysis (See Supplemental Figure 2A, B for an example).

#### LFP Recording

LFP recordings were performed using an open-ephys GUI platform (https://open-ephys.org/) at a sampling rate of 20kHz. Data were acquired with a custom-made headstage using IntanTechnologies’ RHD Electrophysiology Amplifier chip. EEG data and video were recorded simultaneously. Custom designed stimuli trigger generator devices sent triggers directly to the recording system assuring the time precision of the stimuli presentation.

##### Free whisking exploration

Starting 1 week after surgery, animals were habituated daily for 3 days to an open field (15–20 min each time), and baseline activity was recorded while mice were freely exploring the open field. We used a dimly illuminated open field environment (45 by 35 cm), surrounded by 60 cm-high opaque walls and equipped with video monitoring on top of the arena. The open field’s floor and walls were sprayed and wiped clean with 70% ethanol 30 min before the introduction of each animal. On day 4, animals were recorded during free whisking exploration. In particular, two identical textured objects (rougher textured object in Supplemental Figure 1B) were placed in the open field and both LFP and video were recorded for 30 minutes. Exploration bouts were defined by the direct contact between the mouse nose and the objects (see Data Processing/Data Segmentation section below).

##### Whisker stimulation in awake animals

Mice were habituated for 7–10 days to the head-fixed system situated on an air-supported ball treadmill. This custom-made apparatus allowed the animal to run using a larger range of directions than a classical treadmill. The habituation period was progressively longer, starting from 5 up to 25 min daily. We did not notice any difference in the behavior or habituation on the ball among the different experimental groups. Once mice were habituated to staying on the ball for 25 min, they were recorded during 3 minutes of baseline activity and different whisker stimulation protocols. Since mice typically whisk spontaneously at frequencies of 5–15 Hz, and with intermittent bouts lasting 1-4 s (Arakawa & Erzurumlu, 2015), we delivered whisker stimuli at a frequency of either 6 or 10 Hz, with a stimulus duration of 1 s and an interstimulus interval of 2 seconds. In particular, the first protocol consisted of inducing a total of 100 down-up stimulations, each 1s long at 6Hz. The second protocol consisted of 100 stimulations, 1s long at 10Hz. The latter stimulation was chosen because rodents tend to whisk at 10-12Hz for active whisking (Arakawa & Erzurumlu, 2015) and it is consistent with previously published studies (He et al. 2017). Finally, to dissociate intrinsic network dynamics from stimulus-driven frequency entrainment, power spectral density (PSD) analyses were performed using a low-frequency (1 Hz) stimulation protocol. Unlike steady-state paradigms in which oscillatory responses reflect direct entrainment to stimulus frequency, this approach minimizes harmonic contamination and reveals endogenous gamma-band activity associated with circuit-level processing. Each protocol lasted 5 minutes and a period of 2 minutes between each was given to the animals for them to rest. The total duration of each session was 20-25 min.

#### Data Processing

For local field potential (LFP) analysis, EEG signal was down-sampled to 2000 Hz. A notch filter (59.5–60.5 Hz) was applied to remove residual 60 Hz power-line noise contamination. A wideband filter (0.5-150 Hz) was also applied.

##### Data segmentation

Free object exploration periods were automatically detected by using DeepLabCut algorithms. To build the DeepLabCut deep-learning network we manually labelled a variety of contact types: nose-object, body-object, rearing (two frontal paws on the object) and climbing (four paws on the object). Exploration bouts were defined as nose-object interactions that lasted at least 0.4 s. We selected frames in which DeepLabCut detection showed a likelyhood >0.9. We also inspected visually about half of the videos to confirm detection accuracy. Complete immobility periods were calculated using the body center XY coordinates provided by DeepLabCut. In particular, we calculated Euclidean distances of the body center over frames and selected only those periods in which the body center moved less than <1.5 cm/s for at least 1 second. For whisker stimulation analysis we used triggers generated by our custom designed stimuli trigger generator device to segmentate the data into periods of 3000 ms (1000 ms pre- and 2000 ms post-stimulus onset)^76^.

#### Spike Sorting and Unit Classification

Clustering of spikes and unit isolation procedures were performed as described in^77^. Briefly, the power in the 800-9000 Hz range was computed for sliding windows (12.8 ms). Action potentials with a power of > 5 standard deviations (SD) from the baseline mean were selected and their spike features extracted with principal components analysis. Action potentials were then grouped into multiple putative units based on their spike features using an automatic clustering software (http://klustakwik.sourceforge.net). The generated clusters were then manually refined using a graphical cluster-cutting program and only units with clean refractory periods in their auto-correlograms, well-defined cluster boundaries and stability over time were used for further analysis. Putative excitatory neurons (broad-spiking) and PV interneurons (narrow-spiking) were discriminated using the bimodal distribution of spike width (Supplemental Figure 2C).

#### Data Analysis

Signal analysis and quantification was performed using custom MATLAB (The Mathworks Inc., Natik, MA, USA) code, available upon request.

##### Evoked Related Potentials (ERP)

EEG signal was baseline corrected to the mean voltage of the 150 ms prior stimulus onset and averaged over trials. The N1 (baseline-to-peak) component was manually detected from the ERP grand-average waveshape. Analyses were done using Fieldtrip toolbox v 202009.

##### Computation of oscillatory power during free object exploration

Power spectral density (PSD) of LFP signals was estimated using Welch’s method for each exploration epoch. PSDs were then averaged across epochs and normalized by the total spectral power^78^. Logarithmic scaling was used for visualization. Only exploration epochs lasting ≥0.4 s were included to allow reliable estimation of low-frequency oscillations. Immobility periods were defined as epochs in which the body center velocity remained <1.5 cm/s for ≥1 s.

##### Computation of oscillatory power during whisker stimulation

PSD was estimated using Welch’s method for each stimulation epoch (0–1000 ms post-stimulus onset) and averaged across epochs. Spectra were then baseline-corrected using the pre-stimulus period (−500 to 0 ms), following the procedure described in (Carreño-Muñoz et al., 2022), according to the formula:

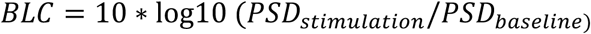

##### Temporal reliability of ERP responses across stimulus repetitions

Temporal reliability of ERP responses was quantified by assessing the consistency of response timing across repeated stimuli within the stimulus train. For each trial and each stimulation, N1 latency was defined as the time of the minimum ERP value within a 20–105 ms post-stimulus window. Analyses were restricted to stimulations 3–10, as differences in ERP timing mainly emerged after the second stimulation. For each stimulation position, latency vectors were constructed containing the latencies measured across all trials. Pairwise Pearson correlations were then computed between latency vectors corresponding to different stimulation positions. Correlation coefficients were Fisher z-transformed to stabilize variance and approximate normality. The median transformed correlation across stimulation pairs was computed for each animal and subsequently back-transformed to Pearson correlation coefficients. This analysis captures the stability of response timing across repeated sensory inputs, providing a measure of how reliably neural circuits maintain temporally precise sensory processing during stimulus trains.

##### Intertrial coherence

Analogous to the phase-locking value, inter-trial coherence allows assessment of the strength of phase coherence across trials in temporal and spectral domains. The inter-trial coherence computation uses only the phase of the complex values given by Morlet’s wavelet transform. Inter-trial coherence measures phase coupling across trials at all latencies and frequencies and is defined by:

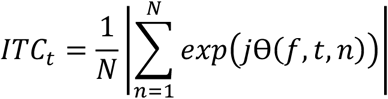

where *j*Θ(*f*, *t*, *n*) represents the phase for a given frequency (f), time point (t) and trial (n). The obtained values are always defined between 0 and 1. Phase-locking values close to 1 indicate strong inter trial phase-locking, thus representing evoked activity while scores closer to 0 indicate a high inter trial phase variability.

##### Quantification of firing rate

The firing rate was computed by dividing the total number of spikes a cell fired in each contact period by the total duration of that period.

##### Spike-field coherence analysis

It was calculated using multitaper methods in order to increase spectral resolution. In this case, we used a chronux function called *coherencycpt*.

### Statistics and reproducibility

Bar graphs represent mean ± SEM. Statistical analyses were performed using Prism 10.0 (GraphPad Software), with the exception of linear mixed model (LMM) analysis that was performed with IBM SPSS V29.0.0. Prior to making comparisons across values, the normality of distribution was tested. Differences between 2 experimental groups were assessed using two-tailed t-test or t-test with Welch’s correction for normally distributed data and Mann-Whitney test for not normally distributed data. Differences between >2 experimental groups were assessed using one-way ANOVA for normally distributed data and one-way ANOVA on Ranks for not normally distributed data. Patch clamp data was analysed using LMM, modelling animal as a random effect and genotype as fixed effect. We used this statistical analysis because we considered the number of mice as independent replicates and the number of cells in each mouse as repeated measures. For the analysis of AP firing, we used two-way ANOVA with Sidak’s multiple comparison *post hoc*.

## DATA AVAILABILITY

Detailed statistic and all data generated and analyzed in the article are available from the corresponding author upon request.

## RESULTS

### PV-cell specific *Tsc1* haploinsufficiency leads to atypical tactile behavior which is rescued by mTORC1 inhibition during the third postnatal week

The rational of this study is based on three observations, 1) sensory processing abnormalities, including tactile disturbances, are often observed in NDDs, particularly in cases where ASD is present as comorbidity^3^; 2) NDDs associated with ASD may share an altered inhibition phenotype, in particular related to PV cell function^9^; 3) abnormalities in mTORC1 signaling have been identified in several NDDs in which ASD is highly prevalent^15^. Whether mTORC1 dysregulation in PV cells is sufficient to induce sensory processing abnormalities, and the underlying mechanisms, are unknown.

To address this question, we used a transgenic mouse carrying a conditional allele of *Tsc1*, which allows cell-specific developmental stage restricted manipulation of Tsc1, crossed to the mouse line with the Cre allele under the control of the PV promoter (PV-Cre^+/−^). This cross generated PV-cell restricted heterozygous mice (*PV-Cre*;*Tsc1^flox/+^*) and their control littermates (*PV-Cre*;*Tsc1^+/+^* and *Tsc1^flox/+^* mice, referred to hereafter as *Tsc1^Ctrl^*). We have previously demonstrated mTORC1 dysregulation in PV cells using this mouse model^17^. First, we asked whether adult mutant mice exhibited alterations in whisker-dependent tactile behavior, by using a texture-specific novel object recognition test (textured NORT, Figure 1A, Supplemental Figure 1)^22–24^. While both conditional heterozygous mice and their control littermates showed a preference for the novel-textured object, mutant mice unexpectedly spent more time exploring it, as reflected by a significantly higher discrimination index (Figure 1B). The total amount of time spent investigating objects during the test did not differ between mutants and control littermates (Figure 1C), indicating that mutant mice did not exhibit an aversion to either object. To investigate whether these behavioral alterations were specific for textured NORT, we tested the novelty seeking behavior of the mice by using a shape NORT in which the objects differed in color and shape but not in texture (Figure 1D-F). In this test, *PV-Cre*;*Tsc1^flox/+^* mice performed similarly to control mice (Figure 1E), suggesting that these mice did not have learning or short-term recognition memory alterations.

**Figure 1.**
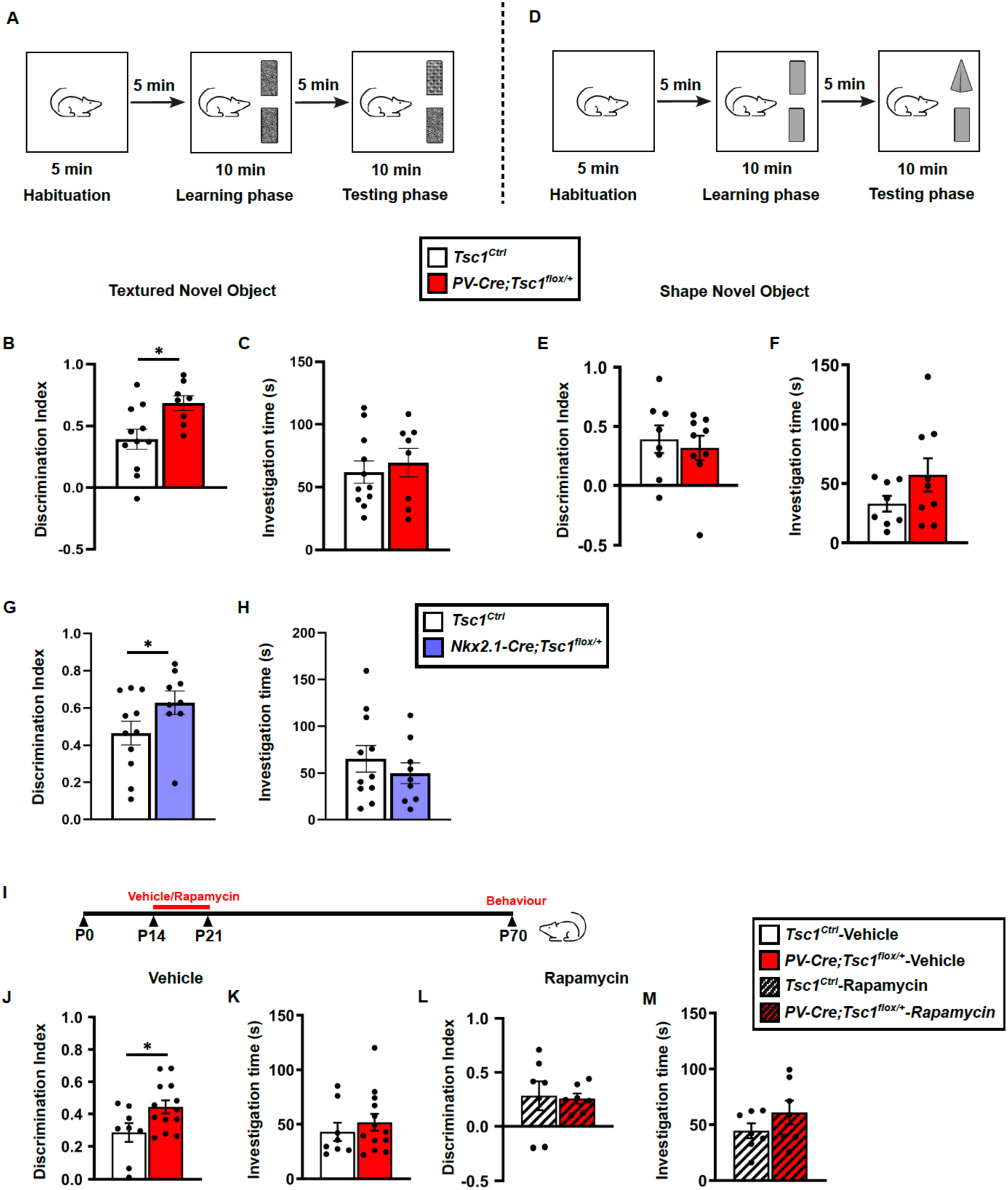
Adult mice with *Tsc1* haploinsufficiency in cortical PV cells show increased preference for the novel textured object. **A,** Schematic representation of texture novel object recognition test. **B,** *PV-Cre;Tsc1^flox/+^* mice show a higher preference for the textured novel object than *Tsc1^Ctrl^*(Welch’s t-test: *p=0.01). **C,** Time spent investigating the objects is similar for both genotypes (Welch’s t-test: p=0.61). Number of mice: *Tsc1*^Ctrl^ n=11; *PV-Cre*;*Tsc1^flox/+^* n=8. **D,** Schematic representation of shape novel object recognition test. **E,** Mutant and control mice show similar shape novel object discrimination index (Welch’s t-test: p=0.6411) and **F,** time spent investigating the objects (Welch’s t-test: p=0.1470). Number of mice: *Tsc1*^Ctrl^ n=8; *PV-Cre*;*Tsc1^flox/+^*n=9. **G,** *Nkx2.1-Cre;TSC1^flox/+^* mice show a higher preference for the textured novel object than *Tsc1^Ctrl^*mice (Mann-Whitney test: *p=0.046), suggesting that *Tsc1* haploinsufficiency restricted to cortical interneurons derived from the medial ganglionic eminence is sufficient to drive increased textured novel object discrimination**. H,** Time spent investigating the objects is similar for both genotypes (Welch’s t-test: p=0.399). Number of mice: *Tsc1*^Ctrl^ n=11; *Nkx2.1-Cre*;*Tsc1^flox/+^*n=9. Bar graph represent mean± SEM. Dots represent individual mice. **I,** Experimental timeline. **J, K,** Discrimination index (**J,** Welch’s t-test: *p=0.0436) and time spent investigating (**K,** Welch’s t-test: p=0.4048) of vehicle-treated mice. Number of mice: *Tsc1*^Ctrl^ n=8; *PV-Cre*;*Tsc1^flox/+^*n=13. **L**, **M,** Rapamycin treatment from P14-21 recues textured novel object discrimination alterations in adult *PV-Cre*;*Tsc1^flox/+^*mice (**L**, Mann-Whitney: p=0.535), without affecting the time spent investigating the objects (**M**, Mann-Whitney: p=0.3176). Number of vehicle-treated mice: *Tsc1*^Ctrl^ n=8; *PV-Cre*;*Tsc1^flox/+^*n=1; number of rapamycin-treated mice: *Tsc1*^Ctrl^ n=7; *PV-Cre*;*Tsc1^flox/+^*n=7.

One caveat is that PV is expressed by subsets of spinal cord neurons and proprioreceptors^25^, thus these neurons might be affected in the mutant mice as well. Since spinal cord PV cells play a critical role in pain gating, we first examined mechanical hypersensitivity via the brush test^26^ and found no significant difference in average nocifensive response score between the two genotypes (0.49±0.9 for *Tsc1*^Ctrl^ vs 0.31±0.9 for *PV-Cre*;*Tsc1^flox/+^* mice, unpaired t-test with Welch’s correction, p=0.15; n=15 *Tsc1*^Ctrl^ and n=14 *PV-Cre*;*Tsc1^flox/+^* mice). To further rule out potential spinal circuit involvement in the observed behavior, we generated *Nkx2-1Cre;Tsc1^flox/+^*mice. Nkx2-1, a transcription factor expressed in interneurons derived from the medial ganglionic eminence (MGE, including PV cells and somatostatin-expressing interneurons), is not detected either in the peripheral nervous system or in the thalamus^27^. Similarly to *PV-Cre*;*Tsc1^flox/+^* mice, *Nkx2-1Cre;Tsc1^flox/+^* mice showed higher discrimination index in the textured NORT compared to control littermates (Figure 1G), suggesting that *Tsc1* haploinsufficiency in MGE-derived interneurons is sufficient to cause increased novel textured object exploration.

We further asked whether this behavioral phenotype was dependent on mTORC1 hyperactivation. We have previously shown that a short-term treatment with the mTORC1 inhibitor, rapamycin, during the third postnatal week prevents the loss of perisomatic synapses formed by PV cells in *PV-Cre*;*Tsc1^flox/+^* mice^17^. We therefore investigated whether this treatment could also rescue alterations in the textured NORT in conditional heterozygous mice. We treated *PV-Cre*;*Tsc1^flox/+^*mice and control littermates daily with either rapamycin (3 mg/kg; i.p.) or vehicle from P14 to P21 and analyzed textured NORT in adult mice (P60-80; Figure 1I). We found that, while vehicle-treated *PV-Cre*;*Tsc1^flox/+^* mice showed significantly higher texture discrimination index than vehicle-treated control littermates (Figure 1J, K), rapamycin-treated *PV-Cre*;*Tsc1^flox/+^* and *Tsc1^Ctrl^* mice showed similar values (Figure 1L, M). Therefore, the third postnatal week represents a critical window during which mTORC1 hyperactivation within PV cells precipitates the emergence of atypical tactile exploratory behavior.

### Transient inhibition of mTORC1 during the third postnatal week prevents the emergence of altered network activity within barrel cortex and prolonged tactile exploration in mutant mice

The effects of PV cell-specific *Tsc1* haploinsufficiency on tactile behavior were unexpected, since most studies reported reduced discrimination in the textured NORT in rodent models of NDDs associated with ASD^4^. Since mutant mice showed increased exploration of the novel textured object compared to control littermates during a 10min exploration period (Figure 1), we wondered whether we could detect further behavioral differences if the mice were allowed to explore the objects longer. At the same time, we sought to characterize potential cortical dysfunctions resulting from *Tsc1* haploinsufficiency in PV cells. Therefore, next we performed extracellular recording of local field potentials (LFP) in adult mutant and control littermates during a 30-minute active exploration of novel textured objects (Figure 2 A, B, Supplemental Figure 2). We defined exploration based on nose-to-object contact, and quantified total exploration time (Figure 2C, D), number of exploration bouts (nose-to-object contacts, Supplemental Figure 3A) and duration of each exploration bout (Supplemental Figure 3B). While mutant and control mice exhibited comparable exploration behavior during the first 15 minutes (Figure 2C, Supplemental Figure 3), mutant mice showed significantly increased total and single-bout exploration time, along with a higher number of exploration bouts, from 15–30 minutes (Figure 2D, Supplemental Figure 3). Notably, total time spent moving did not differ between the two genotypes at any time interval (Figure 2E, F), suggesting that the mutant mouse behavior reflects a specific increase in tactile exploration rather than generalized hyper-locomotion.

**Figure 2.**
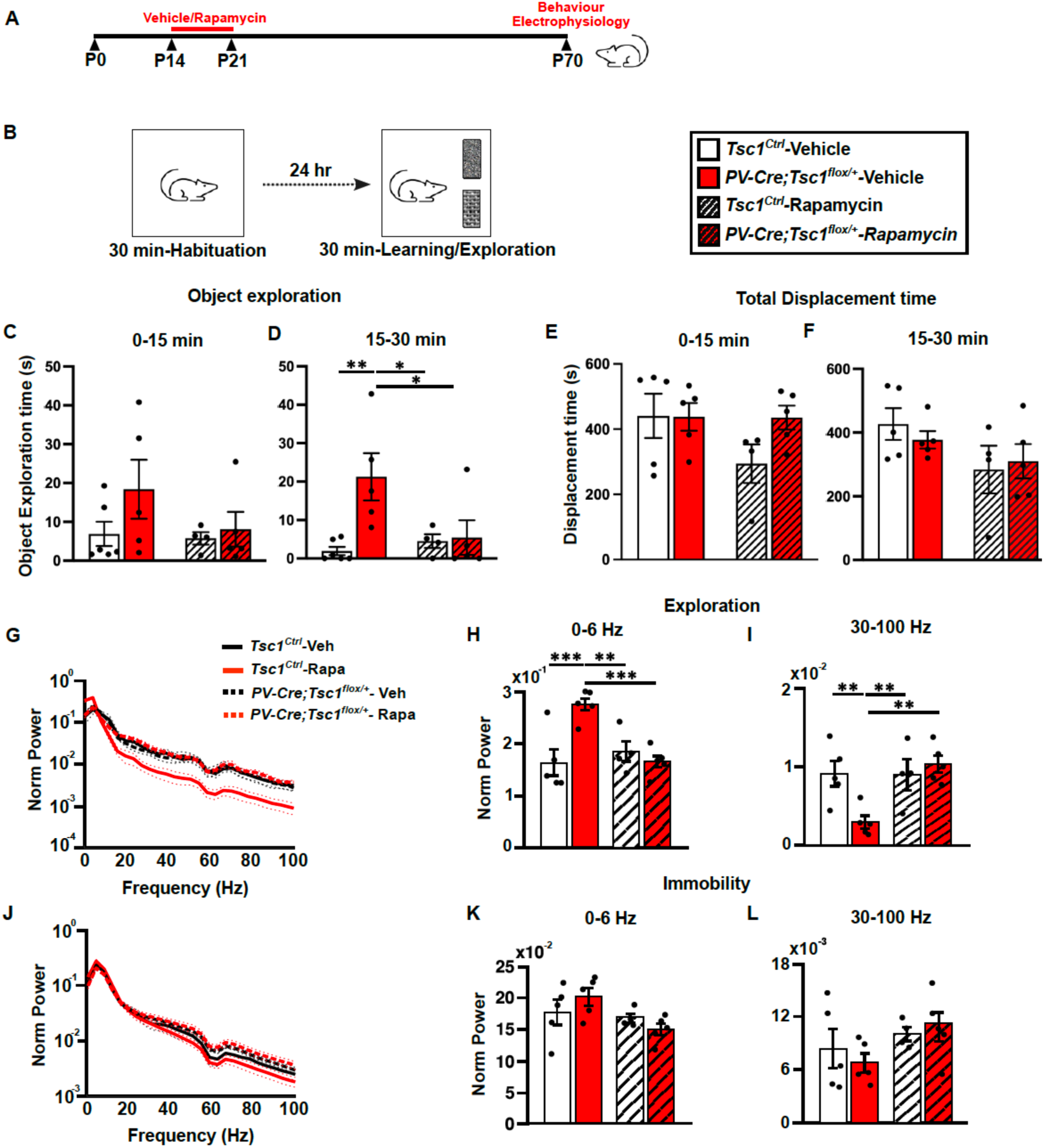
mTORC1 hyperactivity in PV cells during the third postnatal week causes atypical tactile exploratory behavior and state-dependent alterations of network activity within somatosensory circuits. **A,** Experimental timeline. **B,** Experimental approach for in vivo recording during voluntary textured objects exploration. **C, D**, Time spent exploring textured objects in the first (**C,** 0-15min, Two-way ANOVA, interaction p=0.3632) and second half of the exploration period (**D,** 15-30min, Two-way ANOVA, interaction *p=0.0338, Fisher’s LSD multiple comparison, *Tsc1*^Ctrl^-Veh Vs *PV-Cre*;*Tsc1^flox/+^*-Veh **p=0.0022; *PV-Cre*;*Tsc1^flox/+^*-Veh Vs *PV-Cre*;*Tsc1^flox/+^*-Rapa *p=0.0115, *Tsc1*^Ctrl^-Veh vs *PV-Cre*;*Tsc1^flox/+^*-Rapa p=0.5184, *Tsc1*^Ctrl^-Rapa vs *PV-Cre*;*Tsc1^flox/+^*-Veh *p=0.0117). **E**, **F**, Total displacement time between 0-15 (**E,** Two-way ANOVA, interaction P=0.1933) and 15-30 minutes (**F,** Two-way ANOVA, interaction P=0.4774). Displacement is defined as the center of the body moving at a speed > 1.5 cm/s. **G-L**, Power density spectra (**G, J**) and mean power of the 0-6Hz (**H, K**) and gamma band (30-100Hz; **I, L**) recorded in layer 5 of somatosensory cortex during free object exploration (**G**-**I**) and immobility (**J-L**). **H**, **I**, During exploration, LFP power of the 0-6 Hz (**H**) and gamma (**I**) bands is significantly different in vehicle-treated mutant mice compared to both rapamycin-treated mutant and vehicle-treated control mice (**H**, Two-way ANOVA, interaction **p=0.003; Fishers LSD multiple comparison *Tsc1*^Ctrl^-Veh Vs *PV-Cre*;*Tsc1^flox/+^*-Veh ***p=0.0005; *PV-Cre*;*Tsc1^flox/+^*-Veh Vs *PV-Cre*;*Tsc1^flox/+^*-Rap ***p=0.0006; *Tsc1*^Ctrl^-Rap Vs *PV-Cre*;*Tsc1^flox/+^*-Veh **p=0.0044; **I**, Two-way Anova, interaction *p=0.0161; Fishers LSD multiple comparisons test *Tsc1*^Ctrl^-Veh vs *PV-Cre*;*Tsc1^flox/+^*-Veh **p=0.0053; *PV-Cre*;*Tsc1^flox/+^*-Veh Vs *PV-Cre*;*Tsc1^flox/+^*-Rap ** p=0.0014, *Tsc1*^Ctrl^-Rap Vs *PV-Cre*;*Tsc1^flox/+^*-Veh **p=0.0084). **K, L,** During immobility, EEG power at the 0-6 Hz band (**K**) and gamma band (**L**) is unaffected by both genotype and treatment (**K**, Two-way ANOVA, interaction p=0.1517; **L,** Two-way ANOVA, interaction p=0.3832). Number of mice: Vehicle-treated *Tsc1*^Ctrl^ n=5 and *PV-Cre*;*Tsc1^flox/+^* n=5; Rapamycin-treated *Tsc1*^Ctrl^ n=4 and *PV-Cre*;*Tsc1^flox/+^* n=5.

To discern if any potential alteration was state-dependent, we analysed LFP while mice were either actively exploring textured objects or were immobile. Power density spectra analysis revealed a significant alteration in LFP activity in mutant mice, which exhibited a significant power increase in the 0-6 Hz band activity (Figure 2G, H), but a significant power reduction in the broad-band gamma (30-100Hz, Figure 2G, I) during periods of voluntary textured object exploration. In contrast, the activity at all analyzed frequency bands did not show notable differences in mutant vs control mice during complete immobility (Figure 2J-L). Altogether, these data indicate that *Tsc1* haploinsufficiency in PV cells lead to changes in state-dependent network activity within barrel cortex circuits. Of note, mutant adult mice treated with rapamycin during the third postnatal week did not exhibit prolonged textured object exploration (Figure 2D, Supplemental Figure 3) or altered LFP power compared to vehicle- or rapamycin-treated controls (Figure 2G-L). These findings suggest that mTORC1 hyperactivation within PV interneurons during this sensitive developmental window drives the emergence of both tactile-related and network-activity phenotypes.

### Adult mutant mice show reduced temporal precision of whisker-evoked cortical activity, which is rescued by transient inhibition of mTORC1 during the third postnatal week

To elucidate how *Tsc1* haploinsufficiency in PV cells impacts cortical whisker-evoked responses—and to determine if these phenotypes are rescued by short-term rapamycin treatment— we recorded evoked-related potentials (ERPs) following whisker deflection in awake, head-fixed mice. To assess cortical sensory perception, we analyzed ERPs, defined here as LFP activity occurring within 500 ms of stimulus presentation. Since mice typically whisk spontaneously at frequencies of 5–15 Hz, we first analyzed ERP following single 1-sec long stimulation of the whiskers at 6Hz (Figure 3A, B). Compared to control littermates, mutant mice exhibited no difference in amplitude (Figure 3C), but a statistically significant increase in the delay of the N1 component of the ERPs, which was rescued by rapamycin treatment (Figure 3D). We further observed a significant increase of the power of both the evoked 0-6 Hz and broad-band gamma in vehicle-treated *PV-Cre*;*Tsc1^flox/+^* mice compared to vehicle-treated control littermates (Figure 3E-G). Conversely, cortical responses in rapamycin-treated mutant mice did not differ from those recorded in vehicle-treated controls (Figure 3E-G).

**Figure 3.**
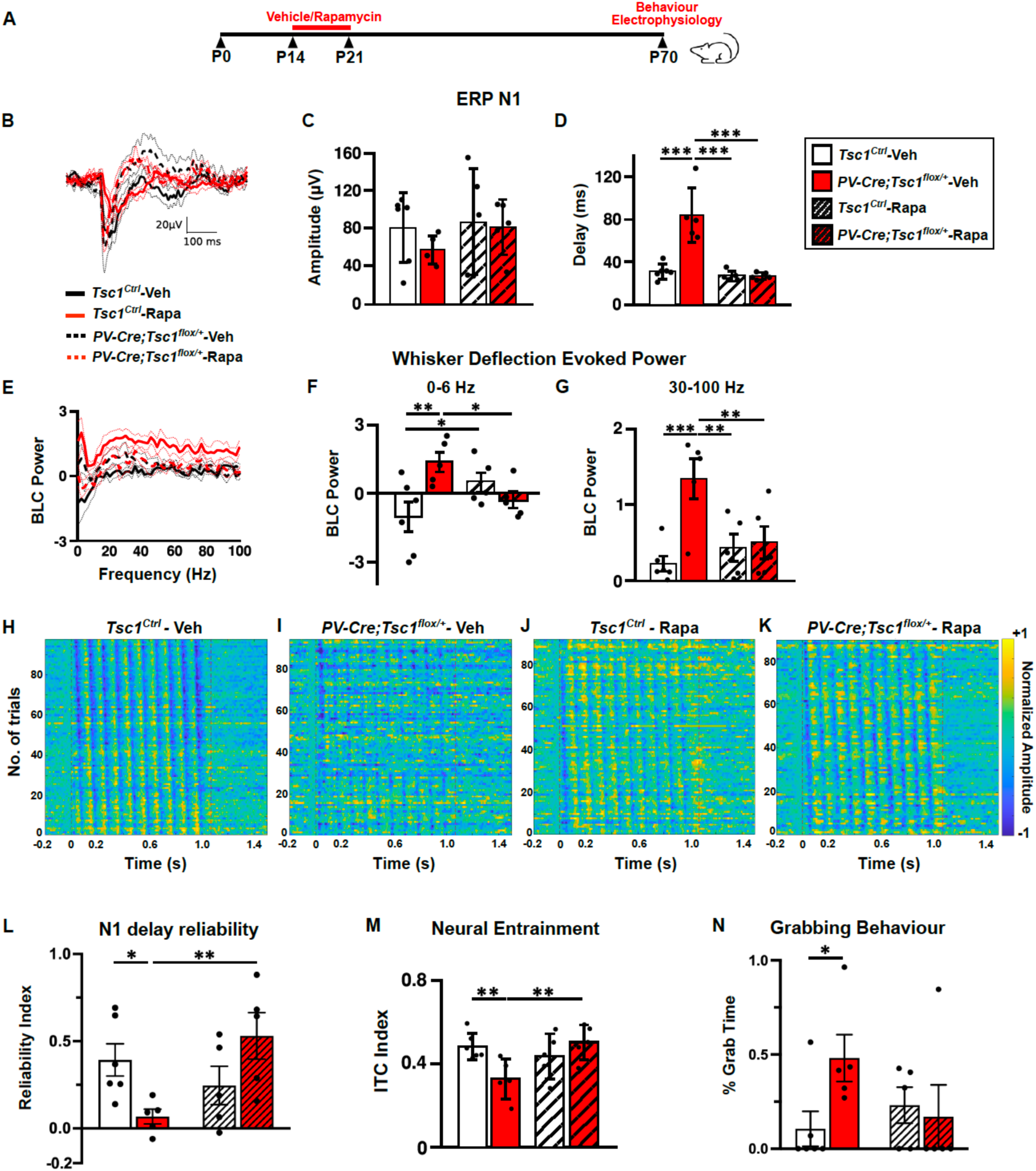
mTORC1 hyperactivity in PV cells during the third postnatal week leads to reduced temporal precision of whisker-evoked cortical activity. **A**, Experimental timeline. **B-D**, Rapamycin treatment rescues ERP delay evoked by whisker deflection in awake head-fixed adult mutant mice. **B**, ERP profile. **C, D,** Bar plots show N1 amplitude (**C)** and delay (**D**). The amplitude is not affected by the genotype or treatment (**C**; Two-way ANOVA, interaction p=0.5633), while the delay is longer in vehicle-treated mutant mice compared to the other 3 experimental groups (**D**, Two-way ANOVA, interaction ***p=0.0003; Fishers LSD multiple comparison test *Tsc1*^Ctrl^-Veh vs *PV-Cre*;*Tsc1^flox/+^*-Veh ***p<0.0001; *Tsc1*^Ctrl^-Rap vs *PV-Cre*;*Tsc1^flox/+^*-Veh ***p<0.0001; *PV-Cre*;*Tsc1^flox/+^*-Veh vs *PV-Cre*;*Tsc1^flox/+^*-Rap *** p<0.0001). **E-G**, Rapamycin treatment rescues gamma power evoked by whisker deflection in awake head-fixed mutant mice. **E**, Baseline corrected (BLC) power density spectra. **F,** Evoked LFP of the 0-6 Hz band is significantly higher in vehicle-treated mutant mice compared to vehicle-treated control mice (Two-way ANOVA, interaction **p=0.0049; Fishers LSD multiple comparison test *Tsc1*^Ctrl^-Veh vs *PV-Cre*;*Tsc1^flox/+^*-Veh **p=0.0027; *Tsc1*^Ctrl^-Rapa vs *Tsc1*^Ctrl^-Veh *p=0.0389; *PV-Cre*;*Tsc1^flox/+^*-Veh vs *PV-Cre*;*Tsc1^flox/+^*-Rapa *p=0.0322). **G**, Evoked LFP power of the gamma band is higher in vehicle-treated mutant mice compared to the other 3 experimental groups (Two-way ANOVA, interaction *p=0.0138; Fishers LSD multiple comparison test *Tsc1*^Ctrl^-Veh vs *PV-Cre*;*Tsc1^flox/+^*-Veh ***p=0.0005; *Tsc1*^Ctrl^-Rapa vs *PV-Cre*;*Tsc1^flox/+^*-Veh **p=0.0041; *PV-Cre*;*Tsc1^flox/+^*-Veh vs *PV-Cre*;*Tsc1^flox/+^*-Rapa **p=0.0073). **H-K,** Examples of trail-by-trail responses evoked by a 10Hz-whisker stimulation vehicle and rapamycin treated control and mutant mice. **L, M**, In mutant mice, rapamycin treatment during the third postnatal week rescues N1 delay reliability (**L**) and entrainment (inter-trial coherence, **M**) following a 10Hz whisker stimulation (**L**, Two-way ANOVA, interaction **p=0.0074; Fishers LSD multiple comparison test *Tsc1*^Ctrl^-Veh vs *PV-Cre*;*Tsc1^flox/+^*-Veh *p=0.0314; *PV-Cre*;*Tsc1^flox/+^*-Veh vs *PV-Cre*;*Tsc1^flox/+^*-Rapa **p=0.0053; **M**, Two-way ANOVA, interaction **p=0.0094; Fishers LSD multiple comparison test *Tsc1*^Ctrl^-Veh vs *PV-Cre*;*Tsc1^flox/+^*-Veh **p=0.0083; *PV-Cre*;*Tsc1^flox/+^*-Veh vs *PV-Cre*;*Tsc1^flox/+^*-Rapa **p=0.0054). **N**, Head-fixed vehicle-treated mutant mice show increased grabbing behavior of the whisker stimulators compared to control littermates. This behavior is not apparent in rapamycin-treated (% of time spent grabbing the pole, Two-way ANOVA, interaction p=0.0911; Fishers LSD multiple comparison test *Tsc1*^Ctrl^-Veh vs *PV-Cre*;*Tsc1^flox/+^*-Veh *p=0.0401). Number of mice: Vehicle-treated *Tsc1*^Ctrl^ n=6 and *PV-Cre*;*Tsc1^flox/+^*n=5; Rapamycin-treated *Tsc1*^Ctrl^ n=5 and *PV-Cre*;*Tsc1^flox/+^*n=5. Bar graphs represent mean ± SEM, dots represent individual mouse data.

When presented with rhythmic sensory stimulations oscillating at a certain frequency, neural activity synchronizes in time with the stimulus, a process referred to as neural entrainment. To explore whether this cortical activity property was affected by *Tsc1* haploinsufficiency in PV cells, we analyzed the inter-trial coherence (ITC) of ERPs following a 10-Hz stimulation train. We chose this stimulation frequency because during object exploration, rodents move their whiskers back and forth rhythmically (whisking) at frequencies around 10-12Hz for active whisking^28^. Trial-by-trial analysis revealed that the N1 response in mutant mice was less precisely time-locked to stimulus onset in vehicle-treated mutant mice compared to vehicle-treated controls, an effect that appeared to be rescued by rapamycin treatment (Figure 3H-K). We next quantified the temporal reliability of ERP responses by assessing N1 delay consistency across repeated stimuli within the train, as well as ITC in both the temporal and spectral domains. Notably, both measures were significantly decreased in mutant compared to control littermates (Figure 3L, M, Supplemental Figure 4A, B), suggesting a reduced temporal precision of whisker-evoked responses in *PV-Cre*;*Tsc1^flox/+^* mice. Furthermore, rapamycin treatment restricted to the third postnatal week was sufficient to prevent neural entrainment deficits in adult mutant mice (Figure 3L, M, Supplemental Figure 4A, C). Altogether, these data show that *Tsc1*-mTORC1 signaling dysregulation in PV cells induced aberrant whisker-evoked activity in adult mice that can be permanently rescued with short-term rapamycin treatment during the third postnatal week.

Of note, while performing these experiments, we noticed that adult mutant mice manifested significantly more grabbing of the whisker stimulator than control littermates (Figure 3N). In particular, 2 out of 6 vehicle-treated control mice tried to grab the pole during whisker stimulation, whereas all vehicle-treated mutant mice did (5 out of 5 mice, Figure 3N). Mutant mice treated with rapamycin manifested notably less grabbing of the stimulator (with only 1 out of 5 mice showing grabbing behavior, Figure 3N), resembling instead the behavior of vehicle-treated control mice. This phenotype, previously reported in a mouse model of Fragile X syndrome (Fmr1 KO mice)^13, 41^, is thought to represent a maladaptive avoidance/defensive behavior^41^. Hence, these data suggest that mTORC1 hyperactivation within PV cells during a sensitive developmental window is sufficient to elicit behavioral manifestations of tactile defensiveness in mice.

### *Tsc1* haploinsufficient PV cells show reduced glutamatergic drive, intrinsic excitability and firing rate in adult mice

We next asked whether and how Tsc1 haploinsufficiency in PV cells affected their input connectivity and function. Since *Tsc1* deletion has been shown to affect glutamatergic synapse formation^29,30^, we quantified glutamatergic synapses density around PV cell somata by immunostaining cortical slices with pre-synaptic (Vglut1-vesicular glutamate transporter 1) and post-synaptic (PSD-95-post-synaptic density 95) markers in cortical layer 5 of adult mice (Figure 4A-D). We found that the density of perisomatic Vglut1+/PSD-95+ puncta was significantly decreased in adult heterozygous mice compared to the controls (Figure 4E). Furthermore, we analyzed the density of asymmetrical synapses (which represent putative glutamatergic inputs) onto PV cell dendrites by electron microscopy (Figure 4F-I). We found that while the density of PV-positive dendrites was not significantly different between *PV-Cre*;*Tsc1^flox/+^*mice and control littermates (Figure 4J), the density of dendrites bearing asymmetrical synapses was significantly reduced in the mutant mice (Figure 4K). To confirm that the loss of glutamatergic input onto PV cells had a functional effect, we performed targeted voltage-clamp recordings of mEPSCs from layer 5 PV interneurons of the barrel cortex of adult *PVcre-Cre*;*RCE*;*Tsc1^+/+^*(control) and *PV-Cre*;*RCE*;*Tsc1^flox/+^* littermates. The RCE allele carried a coding sequence for GFP downstream a floxed stop codon. Using this strategy, we could identify the soma of recombined PV cells thanks to Cre-dependent GFP expression, therefore allowing their targeting recording. We found significantly decreased mEPSC frequency (Figure 4L, M), but unchanged amplitude and charge transfer (Figure 4N, O), in mutant compared to control littermates, suggesting decreased numbers of functional excitatory synapses onto PV cells, in agreement with our anatomical data.

**Figure 4.**
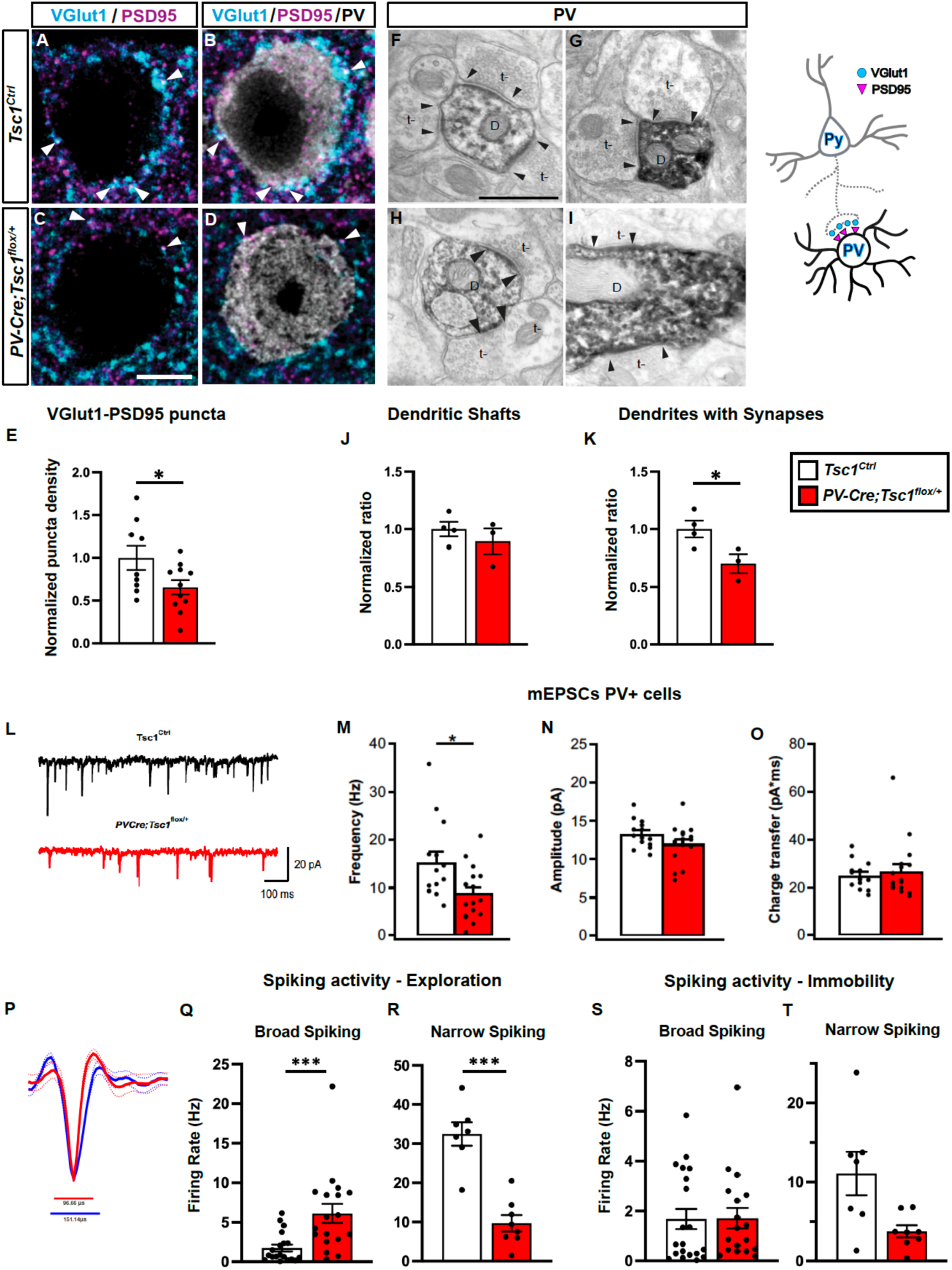
Adult *Tsc1* haploinsufficient PV cells have reduced excitatory drive. **A-D,** Representative confocal images of somatosensory cortex coronal sections immunostained for Vglut1 (cyan), PSD-95 (magenta) and PV (gray) from adult *Tsc1*^Ctrl^ (**A, B**) and *PV-Cre*;*Tsc1^flox/+^* (**C, D**) mice. Scale bar: 5µm. White arrowheads indicate Vglut1/PSD-95 colocalized boutons. **E**, Vglut1/PSD-95 colocalized puncta onto layer 5 PV cell somata are significantly decreased in *PV-Cre*;*Tsc1^flox/+^* mice (Welch’s t-test: *p=0.0417). Number of mice: n=9 for *Tsc1*^Ctrl^ and n=11 for *PV-Cre;Tsc1^flox/+^*. **F-I,** PV-immunolabeled dendritic branches in somatosensory cortex of adult *Tsc1*^Ctrl^ (**F**, **G**) and *PV-Cre;Tsc1^flox/+^* (**H**, **I**) showing multiple asymmetric synapses (arrowheads) between unlabeled axon terminal (t-) and a labeled dendritic shaft (indicated with a D in the microphotographs). Scale bar, 500 nm. **J,** Overall, PV+ dendritic branch density is no different between the two genotypes (Welch’s t-test, p=0.4665). **K,** Quantification of PV+ dendrites bearing synapses shows a significant decrease in *PV-Cre;Tsc1^flox/+^* mice compared to control littermates (Welch’s t-test, *p=0.0422). Arrowheads indicate presence of PSDs (post synaptic density) in the dendrites. Number of mice: *Tsc1^Ctrl^*, n=4; *PV-Cre*;*Tsc1^flox/+^*, n=3. Bar graphs represent mean ± SEM. Dots represent single mouse data. **L**, Representative traces of mEPSCs recorded in PV cells from P60-P70 *Tsc1*^Ctrl^ (black trace) and *PV-Cre;Tsc1^flox/+^* mice (red trace). **M**, mEPSC frequency is significantly decreased in *PV-Cre;Tsc1^flox/+^* mice compared to control littermates (Linear Mixed Modeling (LMM), *p=0.041). **N, O,** mEPSC amplitude (**N)** and charge transfer (**O**) are not different between genotypes (LMM, p>0.05 for both). *Tsc1^Ctrl^*, n=14 cells from 3 mice. *PV-Cre*;*Tsc1^flox/+^*, n=16 cells from 3 mice. Bar graphs represent mean ± SEM. Dots represent individual neurons. **P,** Average spike waveform normalized to the maximal height for the two groups of neuron types, broad-spiking (blue) and narrow-spiking (red, putative PV interneurons), included in the analysis. Dotted lines represent standard deviation. Red and blue lines show the mean width of the corresponding cell type. **Q, R,** Average firing rate of broad-spiking (**Q**) and narrow-spiking (**R,** putative PV cells) neurons from layer 5 of somatosensory cortex recorded during free object exploration. In mutant mice, broad-spiking neurons show increased firing rate (**Q**, LMM: ***p=0.001), while narrow-spiking neurons show decreased firing rate compared to control littermates (**R**, LMM: ***p<0.001) during object exploration. **S**, **T**, Average firing rate of broad-spiking (**S**) and narrow-spiking (**T**) neurons during immobility. There is no significant difference between the genotypes (**S**, LMM: p=0.973; **T**, LMM: p=0.066). **Q-T:** Number of cell: N= 20 broad-spiking and 7 narrow-spiking from 5 *Tsc1*^Ctrl^ mice; N= 18 broad-spiking and 8 narrow-spiking from 5 *PV-Cre*;*Tsc1^flox/+^* mice. Bar graph represent mean± SEM. Dots represent individual cells.

To assess PV cell intrinsic excitability, we performed whole-cell current-clamp recordings, identified rheobase and then recorded F-I curves at current levels above rheobase. Whereas passive and active membrane properties were not significantly different between the two genotypes (Supplemental Figure 5A), mutant PV interneurons fired significantly fewer action potentials in response to the same depolarizing current injection when compared to control mice (Supplemental Figure 5B). Altogether, these results show that *Tsc1*-haploinsufficiency in PV cells altered their functional maturation, by reducing both their glutamatergic inputs and intrinsic excitability.

To assess whether these alterations translated in changes of PV cell firing *in vivo*, we performed extracellular recording of single units in layer 5 of the barrel cortex in freely moving adult mutant and control littermates. Single units were classified as broad-spiking (putative excitatory neurons) and narrow-spiking (putative PV interneurons) based on waveform^31,32^ (Figure 4P). As described above, mice were placed in an open field where they were free to explore two novel identical textured objects for 30’. Consistently with the observed power spectral changes in the LFP at 0-6 Hz and broad-band gamma (Figure 2G-I), the spontaneous firing rate of broad-spiking neurons was significantly increased (Figure 4Q), while that of narrow-spiking neurons was decreased (Figure 4R) in mutant compared to control mice during object investigation. This data suggest that the activity of pyramidal neurons is increased by the disinhibition of PV neurons during voluntary object exploration; however, we cannot exclude that other neuron types, for example somatostatin-expressing interneurons, are included in the broad spiking neuronal population. Of note, while no statistical differences in the spontaneous firing rate of either neuronal population were detected during immobility (Figure 4S, T), the spontaneous firing rate of narrow-spiking neurons showed a trend toward reduced firing rate in mutant mice (Figure 4T, LMM, p=0.066), consistent with the reduced excitatory drive observed *ex-vivo*.

Altogether, these data indicate that *Tsc1* haploinsufficiency in PV cells leads to their hypoactivity within barrel cortex circuits.

### *Tsc1* haploinsufficient cortical PV cells lose glutamatergic input after the third postnatal week

Since glutamatergic synapse formation onto PV cells occurs during the first postnatal weeks^33^, we next asked whether *Tsc1* haploinsufficiency affected the development of Vglut1+/PSD95+ putative synapses onto PV cells in preadolescent mice (P22, Supplemental Figure 6A-D). As Cre expression in the PV-Cre line only reaches a plateau by the 4^th^ postnatal week, we focussed our analysis on GFP+ neurons in *PV-Cre;RCE;Tsc1^+/+^* (*Tsc1*^Ctrl^) and *PV-Cre;RCE;Tsc1^flox/+^* mice. In contrast to what we observed in adult mice, Vglut1+/PSD-95+ density was not significantly different in preadolescent conditional heterozygous mice compared to control littermates (Supplemental Figure 6E). These data suggest that glutamatergic synapse formation onto *Tsc1* haploinsufficient PV cells occurs normally during the first 3 postnatal weeks; however, these synapses are either subsequently lost or fail to increase in density in mutant mice. The formation of excitatory synapses onto PV cells requires a complex molecular program of synaptic proteins, including adhesion molecules^33^. We next analysed the expression of two of these adhesion molecules, SynCAM1 and Neuroligin 3 (Nlg3), in preadolescent *PV-Cre;RCE;Tsc1^flox/+^* mice compared to control littermates. SynCAM1 expression levels were significantly reduced in layer 5 PV cell dendrites of mutant mice compared to control littermates (Supplemental Figure 7A-E). This was not due to a general reduction of SynCAM1 expression levels, since we did not detect any significant difference in the neuropile between the two genotypes (Supplemental Figure 7E). Similarly, the density of Nlgn3 puncta localized with PSD95 onto layer 5 PV cell somata was reduced in mutant mice (Supplemental Figure 7F-J). Conversely, Nlgn3/PSD95 puncta density onto pyramidal cell bodies was not affected (Supplemental Figure 7J). Thus, it is possible that the downregulation of these two molecules in the first postnatal month may be a first step contributing to the reduced density of excitatory inputs on PV cells observed in adult *Tsc1* haploinsufficient mice.

### Short-term whisker trimming during the third postnatal week dampens down pS6 hyperactivation and prevents the development of PV cell connectivity deficits in *PV-Cre*; *Tsc1^flox/+^* mice

mTORC1 is a signaling hub that regulates neuronal metabolism in response to multiple environmental cues, including plasticity-inducing neuronal activity patterns^34,35^ and activity-dependent neurotrophic factors (for example, BDNF^36^). We thus asked whether mTORC1 hyperactivation due to *Tsc1* haploinsufficiency might be dampened down by reducing sensory inputs. To directly test this hypothesis, we transiently restricted neonatal vibrissal inputs by cutting the whiskers close to the intersection with the skin. Whiskers of *PV-Cre*;*RCE*;*Tsc1^flox/+^* mice and their control littermates (*PV-Cre*;*RCE*;*Tsc1^+/+^*) were trimmed every two days to within 1 mm of the skin from P14 to P21. We chose this time window for several reasons: 1) it coincides with the maturation of complex PV cell innervation, a process highly modulated by neuronal activity and sensory experience^19,37^, 2) pS6 expression, one of the direct downstream effectors of mTORC1, significantly increases during this developmental time window in PV, but not glutamatergic, neurons in the somatosensory cortex^17^, 3) active exploratory whisking emerges at around P14 in mice^28^, at a time that coincides with network activity reaching its final mature stage^38,39^. pS6 expression is a well-established proxy for mTORC1 hyperactivation^29,40^. As expected, pS6 intensity levels were significantly higher in conditional mutant preadolescent mice compared to control littermates (Figure 5A-Q). In contrast, there was no significant difference between whisker-deprived mutant and sham control mice (Figure 5Q), suggesting that limiting sensory experience by whisker deprivation was sufficient to dampen down mTORC1 hyperactivation caused by *Tsc1* haploinsufficiency.

**Figure 5.**
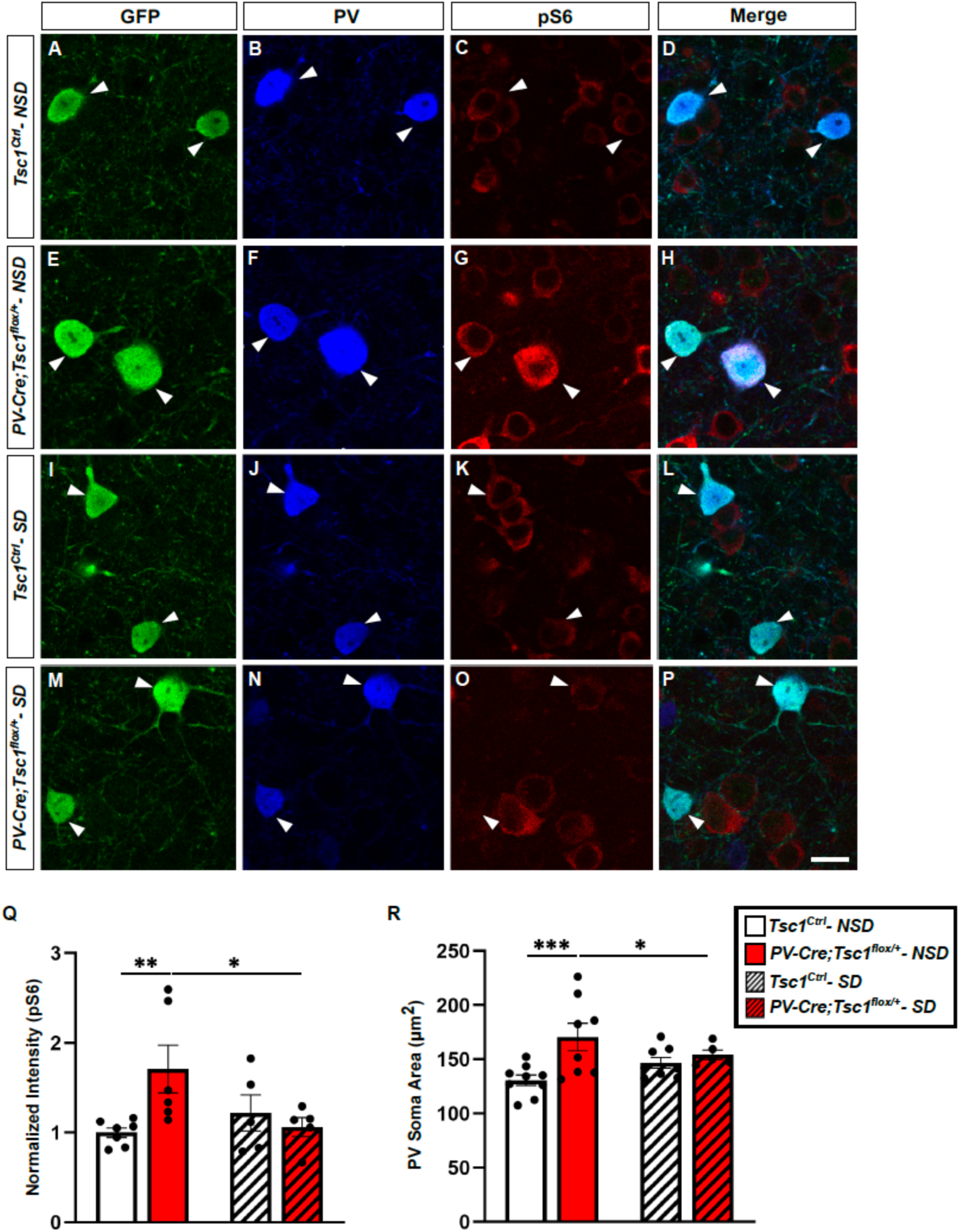
Whisker deprivation during the third postnatal week dampens down pS6 upregulation and prevents soma size hypertrophy in mutant mice. **A-P,** Representative immunostained sections of somatosensory cortex labeled for GFP (green), PV (blue) and pS6 (red) in P22 sham (non-sensory deprived mice, NSD) *Tsc1*^Ctrl^ (**A-D**) and *PV-Cre;Tsc1^flox/+^* (**E-H**) mice, and in sensory deprived mice (SD) *Tsc1*^Ctrl^ (**I-L**) and *PV-Cre;Tsc1^flox/+^* (**M-P**) mice. Scale bar: 15 μm. Arrowheads show GFP/PV/pS6 colocalized somata. **Q**, At P22, pS6 intensity is significantly increased in sham mutant mice compared to sham control mice, while it is not significantly different in sensory deprived mutant mice compared to sham control mice (Two-way ANOVA, interaction *p=0.0224, Fisher’s LSD multiple comparison test: *Tsc1*^Ctrl^**-** NSD vs *PV-Cre;Tsc1^flox/+^*- NSD, **p=0.0062; *PV-Cre;Tsc1^flox/+^*- NSD vs *PV-Cre;Tsc1^flox/+^*- SD, *p=0.0185). Number of mice: n=7 for *Tsc1^Ctrl^*- NSD, n=6 for *PV-Cre;Tsc1^flox/+^*- NSD, n=5 for *Tsc1^Ctrl^*- SD, n=5 for *PV-Cre;Tsc1^flox/+^*- SD. **R**, PV cell soma size is larger in sham adult mutant mice compared to the control mouse groups (Two-way ANOVA, interaction p=0.058, Fisher’s LSD multiple comparison test *Tsc1*^Ctrl^**-** NSD vs *PV-Cre;Tsc1^flox/+^*- NSD ***p=0.0009; *Tsc1*^Ctrl^**-** NSD vs *PV-Cre;Tsc1^flox/+^*- SD p=0.0652; *PV-Cre;Tsc1^flox/+^*- NSD vs *Tsc1*^Ctrl^**–** SD, *p=0.0378). Number of mice: n=8 for *Tsc1^Ctrl^*- NSD, n=8 for *PV-Cre;Tsc1^flox/+^*- NSD, n=7 for *Tsc1^Ctrl^-* SD, n=5 for *PV-Cre;Tsc1^flox/+^*- SD. Bar graphs represent mean ± SEM. Dots represent individual neurons.

Based on this observation, we next asked whether this time-restricted manipulation had long-term effects of PV cell connectivity. First, we analyzed PV cell soma size since somatic hypertrophy has been consistently reported as an effect of mTORC1 hyperactivity in different cell types^17,29,40^. Our analysis showed that PV soma area was significantly larger in sham, but not whisker-deprived, adult mutant mice compared to sham controls (Figure 5R). We then analyzed PV cell glutamatergic inputs, intrinsic excitability and output connectivity. Whisker-deprivation prevented the loss of glutamatergic inputs onto PV cells, analyzed by immunolabeling of synaptic markers (Vglut1+/PSD-95+ puncta, Figure 6A-E) and mEPSC recording (Figure 6F-I). Of note, in adult mutant mice that were whisker-deprived during the 3^rd^ postnatal week, PV interneurons fired more action potentials in response to the same depolarizing current injection when compared to PV cells recorded from sham mutant mice (Supplemental Figure 8A, B). In contrast, PV cells in whisker-deprived control mice showed a strikingly reduction in firing rate compared to PV cell recorded from sham control mice (Supplemental Figure 8C, D). A recent study showed that 1-day whisker deprivation between P18-P23 caused a rapid and significant reduction of layer 2/3 PV cell intrinsic excitability, which was however absent in *Tsc2^+/-^* mice^18^. Our data add to this study by showing that one week-long whisker deprivation causes long-term alterations of layer 5 PV cell excitability, which are drastically different in controls compared to PV-specific *Tsc1* haploinsufficient mice.

**Figure 6.**
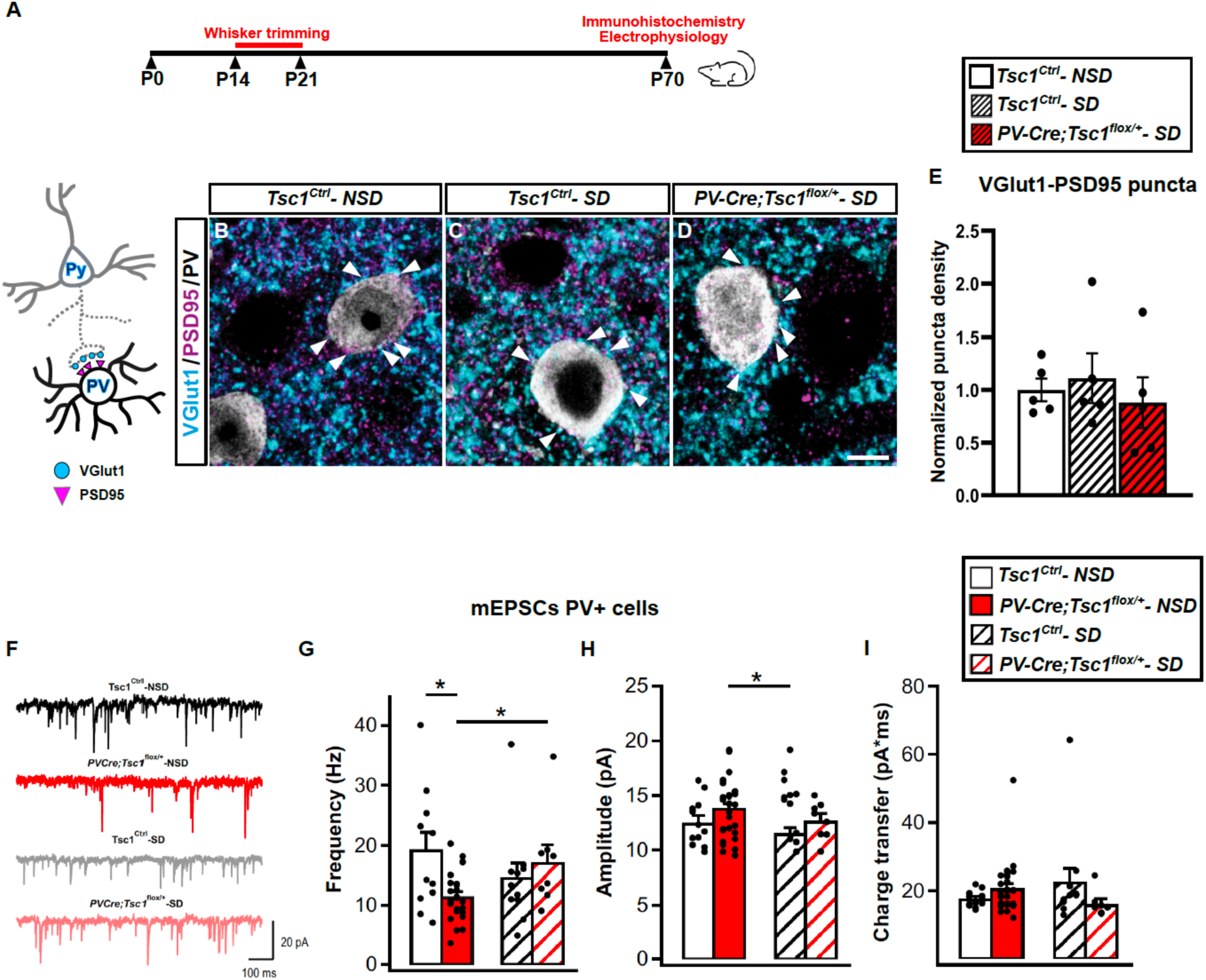
Whisker deprivation during the third postnatal week prevents loss of PV cell glutamatergic inputs in adult mutant mice. **A**, Schematic for sensory deprivation paradigm. **B-D,** Representative immunostained sections of somatosensory cortex labeled for Vglut1 (cyan), PSD-95 (magenta) and PV (gray) in P70 *Tsc1*^Ctrl^ non-deprived mice (NSD) (**B**), *Tsc1*^Ctrl^ deprived mice (SD) (**C**) and *PV-Cre;Tsc1^flox/+^*- SD (**D**) mice. Scale bar: 5μm. Arrowheads show Vglut1/PSD-95 colocalized boutons. **E**, Bar graph shows the density of Vglut1/PSD-95 colocalized puncta impinging onto layer 5 PV cell somata (One-way ANOVA, p=0.7299). Number of mice: n=5 for each experimental group. **F**, Representative traces of mEPSCs recorded in PV cells from adult *Tsc1^Ctrl^* – NSD (black trace), *PV-Cre;Tsc1^flox/+^*- NSD (red trace), *Tsc1^Ctrl^*- SD (gray trace) and *PV-Cre;Tsc1^flox/+^*- SD (pink trace). **G-I**, Summary bar graphs for mEPSC frequency (**G**), amplitude (**H**) and charge transfer (**I**). **G,** mEPSC frequency is significantly reduced in sham (non-sensory deprived, NSD) mutant (*PV-Cre;Tsc1^flox/+^*- NSD) compared to both sham control mice (*Tsc1^Ctrl^*- NSD, Linear Mixed Modeling (LMM), *p=0.006) and sensory-deprived (SD) mutant mice (*PV-Cre;Tsc1^flox/+^* - SD, LMM, *p=0.011). (**H**) mEPSC amplitude in sham mutant mice is significantly higher than in sensory deprived control mice (LMM, *p=0.030). **I**, Charge transfer is not different between the 4 experimental groups (LMM, p>0.05). *Tsc1^Ctrl^*- NSD, n=12 cells from 3 mice. *PV-Cre;Tsc1^flox/+^*- NSD, n=27 cells from 6 mice. *Tsc1^Ctrl^*- SD, n=12 cells from 3 mice. *PV-Cre;Tsc1^flox/+^*- SD, n=8 cells from 3 mice. Bar graphs represent mean ± SEM. Dots represent individual neurons.

To analyze PV cell output connectivity, we quantified perisomatic PV cell synapse density around layer 5 pyramidal neurons by immunostaining cortical slices with presynaptic (PV) and postsynaptic (gephyrin) markers (Figure 7A,B-E, F-I). *Tsc1* haploinsufficiet PV cells formed less GABAergic synapses onto pyramidal cell somata in adult mice (Figure 7F-I, K), confirming our previous data^17^. Of note, sensory deprivation during the third postnatal week was sufficient to rescue the density of PV+/Gephyrin+ putative synapses onto target cell somata in adult mutant mice (Figure 7K). In contrast to what was observed in adult mice, preadolescent (P22) mutant mice showed excessive numbers of perisomatic boutons formed by PV cells around layer 5 pyramidal cell somata (Figure 7B-E, J), consistently with a previous report^17^. Whisker-deprivation during the third postnatal week was sufficient to prevent the premature formation of these putative synapses in preadolescent mice (Figure 7J).

**Figure 7.**
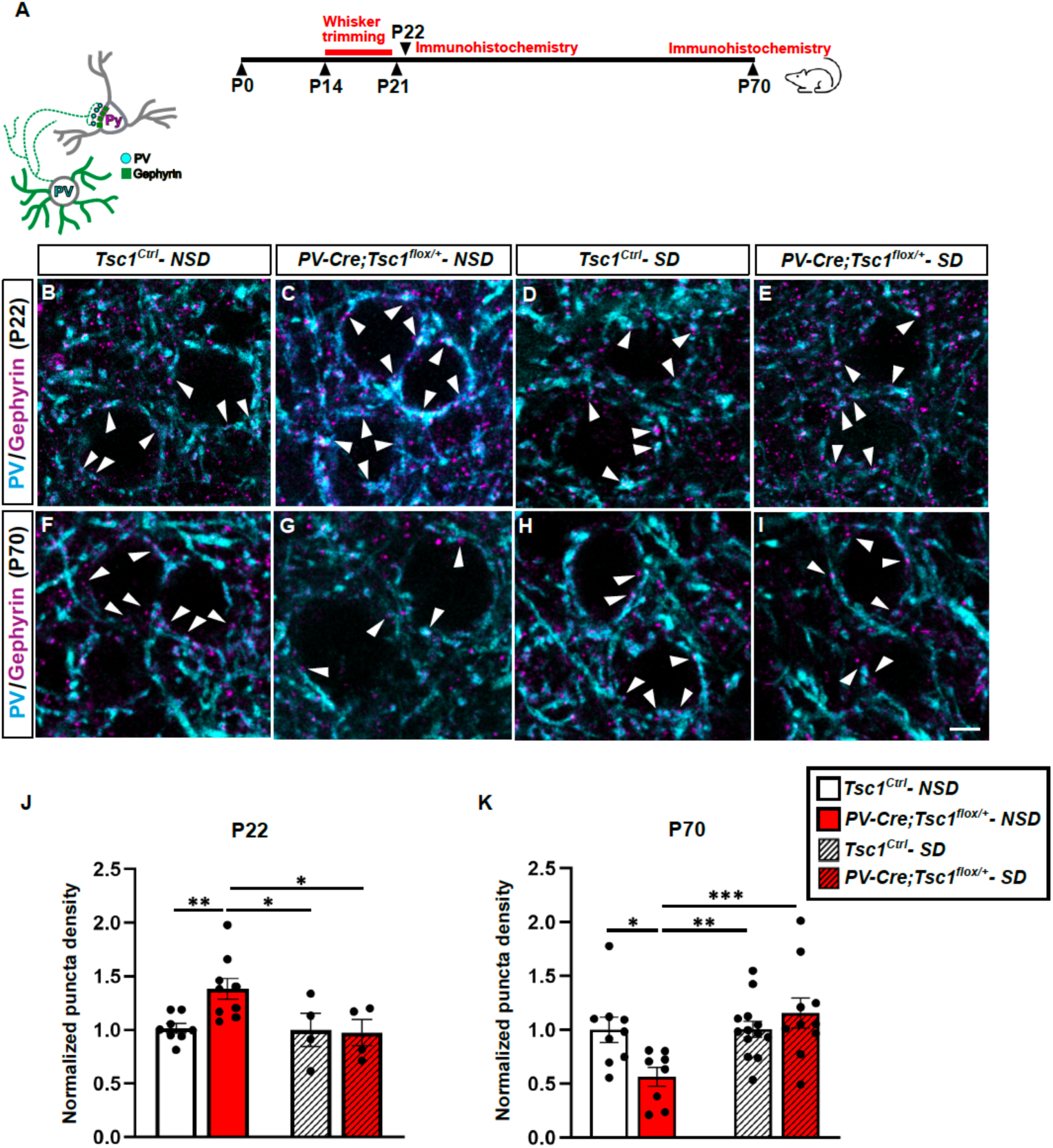
Whisker deprivation during the third postnatal week normalizes the developmental trajectory of PV cell output connectivity in mutant mice. **A**, Schematic of experimental approach. **B-I,** Representative immunostained sections of somatosensory cortex labeled for PV (cyan) and Gephyrin (magenta) in preadolescent (P22, **B-E**) and adult (∼P70, **F-I**) naïve (no sensory deprivation, NSD) *Tsc1*^Ctrl^ (**B, F**) and *PV-Cre;Tsc1^flox/+^* mice (**C, G**), and in sensory-deprived (SD) *Tsc1*^Ctrl^ (**D, H**) and *PV-Cre;Tsc1^flox/+^*(**E, I**) mice. Scale bar: 5μm. Arrowheads indicate PV+Gephyrin+ puncta. **J,** PV/Gephyrin colocalized puncta density is significantly higher in pre-adolescent sham *PV-Cre;Tsc1^flox/+^* than in sham *Tsc1^Ctrl^*mice, but this phenotype is rescued by whisker deprivation between P14-P21 (Two-way ANOVA, interaction p=0.0772, Fisher’s LSD multiple comparison test: *Tsc1*^Ctrl^**-** NSD vs *PV-Cre;Tsc1^flox/+^*- NSD, **p=0.0057; *PV-Cre;Tsc1^flox/+^*- NSD vs *PV-Cre;Tsc1^flox/+^*- SD, *p=0.0114; *PV-Cre;Tsc1^flox/+^*- NSD vs *Tsc1*^Ctrl^- SD, *p=0.0164). Number of P22 mice: n=8 for *Tsc1*^Ctrl^- NSD, n=9 for *PV-Cre;Tsc1^flox/+^*- NSD; n=4 for *Tsc1*^Ctrl^- SD, n=4 for *PV-Cre;Tsc1^flox/+^*- SD. **K,** PV/Gephyrin colocalized puncta density is significantly lower in adult sham *PV-Cre;Tsc1^flox/+^*than in *Tsc1^Ctrl^* mice, but the synaptic loss is prevented by whisker deprivation between P14-P21 (Two-way ANOVA, interaction **p=0.01, Fisher’s LSD multiple comparison test *Tsc1*^Ctrl^**-** NSD vs *PV-Cre;Tsc1^flox/+^*- NSD, *p=0.011; *PV-Cre;Tsc1^flox/+^*- NSD vs *PV-Cre;Tsc1^flox/+^*- SD, ***p=0.0007; *PV-Cre;Tsc1^flox/+^*- NSD vs *Tsc1*^Ctrl^ **–** SD, **p=0.0057). Number of adult mice: n=9 for *Tsc1*^Ctrl^- NSD, n=8 for *PV-Cre;Tsc1^flox/+^*- NSD; n=13 for *Tsc1*^Ctrl^- SD, n=10 for *PV-Cre;Tsc1^flox/+^*- SD. Bar graphs represent mean ± SEM. Dots represent single mouse data.

Taken together, these data suggest that short-term sensory deprivation during a sensitive postnatal window is sufficient to prevent deficits of PV cell excitatory input and output connectivity caused by mTORC1 hyperactivation.

### Short-term whisker trimming during the third postnatal week prevents atypical tactile behavior and aberrant whisker-evoked cortical activity in adult *PV-Cre*; *Tsc1^flox/+^* mice

Adult sensory perception crucially depends on sensory experience during critical postnatal periods (Reh et al. 2020). In *PVCre;Tsc1^flox/+^* adult mice, PV cell excitatory synaptic inputs (Figure 4A-M), firing rate (Figure 4R, T, Supplemental Figure 5) and output connectivity (Figure 7) are significantly reduced, pointing to a PV cell hypoactivity phenotype. The imbalanced activation of PV cells versus pyramidal neurons might be in part responsible for changing the developmental trajectory of neuronal circuits within sensory cortices, thereby undermining the development of typical sensory processing. Since whisker deprivation restricted to the third postnatal week was sufficient to prevent long-term alterations in PV cell soma size (Figure 5), glutamatergic inputs (Figure 6) and output connectivity (Figure 7) in mutant mice, we tested whether this short-term sensory manipulation had long-term effects on both cortical activity within barrel cortex circuits and tactile behaviors.

In the textured NORT, whisker-deprived mutant and control littermates did not show enhanced preference for the novel textured object compared to whisker-deprived control littermates (Figure 8A-D). We next recorded LFP, while mice were free to explore two textured objects for 30 min. Whisker-deprived mutant and control mice did not show differences in their spontaneous tactile exploratory behavior, which resembled that of control mice (compare Figure 2 C, D and Supplemental Figure 3 with Figure 8E-I and Supplemental Figure 9). We also did not detect differences in locomotion (Figure 8H, I). Further, LFP recording showed no genotype-related difference in oscillation power either during voluntary object exploration or immobility (Figure 8J-M). In head-fixed awake mice, we did not detect any genotype-related difference in ERP dynamics (Figure 8N-P) and evoked oscillation power (Figure 8Q, R). Finally, the temporal reliability and inter-trial coherence of ERPs evoked by a 10-Hz stimulation train in conditional heterozygous adult mice was similar to the levels observed in both non-deprived and deprived control mice (Figure 8S-V, Supplemental Figure 10, compare with Figures 3L, M). Finally, adult *PV-Cre*; *Tsc1^flox/+^* mice which were whisker-deprived during the third postnatal week spent less time grabbing the stimulator compared to non-deprived mutant mice (0.16±0.07% in 4 whisker-deprived vs 0.48±0.12% in 5 non-deprived mutant mice, Mann Whitney p=0.025). Conversely, grab time was not significant different between whisker-deprived *PV-Cre*; *Tsc1^flox/+^* and non-deprived control mice (0.16±0.07% in 4 whisker-deprived mutant vs 0.11±0.17% in 6 non-deprived control mice, Mann Whitney p=0.16).

**Figure 8.**
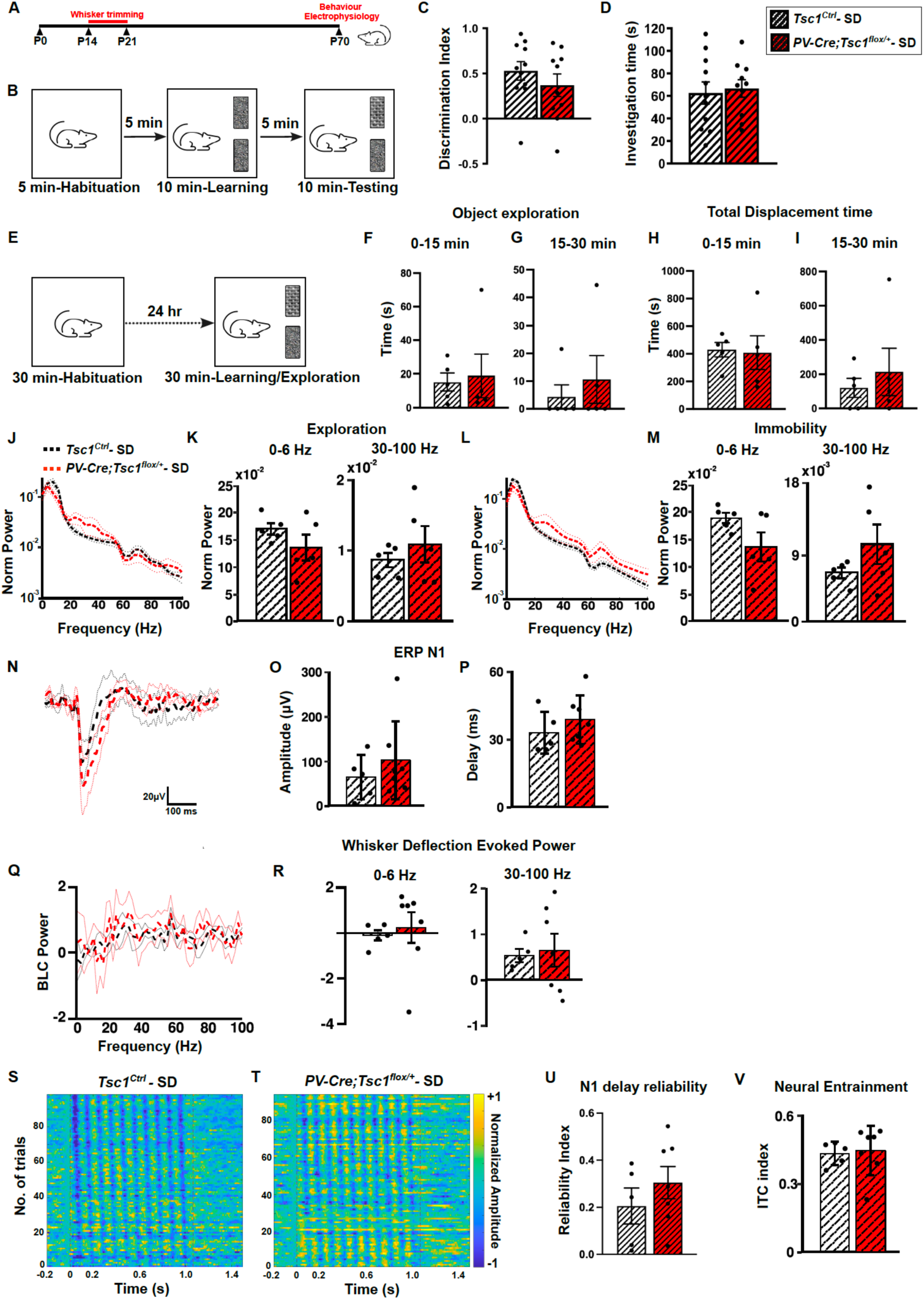
Whisker deprivation during the third postnatal week prevents the development of atypical textured object exploration and cortical activity in adult mutant mice. **A**, Schematic for sensory deprivation paradigm. **B,** Experimental approach for the texture discrimination test. **C, D,** Sensory deprivation between P14-21 rescues texture discrimination in adult mice (**C**, Welch’s t-test, p=0.3363). **D,** time spent investigating the objects (Welch’s t-test: p=0.7538). Number of mice: n=11 for *Tsc1*^Ctrl^ and n=10 for *PV-Cre;RCE;Tsc1^flox/+^*. **E**, Experimental approach for in vivo recording during voluntary textured objects exploration. **F, G**, Time spent exploring textured objects in the first (**F,** 0-15min, Mann-Whitney test, p=0.69) and second half of the exploration period (**G,** 15-30min, Mann-Whitney test, p=0.61). **H**, **I**, Total displacement time between 0-15 (**H,** Mann-Whitney test, p=0.42) and 15-30 minutes (**I**, Mann-Whitney test, p>0.999). **J-M**, Power density spectra (**J, L**) and mean power of the 0-6 Hz and gamma band (**K, M**) recorded in layer 5 of somatosensory cortex during free object exploration (**K**, 0-6Hz, Welch’s t-test, p=0.2179; 30-100Hz, Welch’s t-test, p=0.4245) and immobility (**M,** 0-6 Hz, Welch’s t-test, p=0.0985; 30-100Hz, Welch’s t-test, p=0.1939). LFP power during exploration is rescued in sensory deprived mutant mice. **N-P**, Sensory deprivation rescues ERP delay and gamma power evoked by whisker deflection in awake mutant mice. **N**, ERP profile**, O,P,** Bar plots show N1 amplitude (**O**, Mann-Whitney test, p=0.4168) and delay (**P**, Mann-Whitney test, p=0.3735). **Q**, Baseline corrected (BLC) Power Density Spectra. **R,** Bar plots showing BLC evoked power at 0-6 Hz (Mann-Whitney test, p=0.7011) and gamma (30-100 Hz; Mann-Whitney test, p=0.7978). Number of sensory-deprived mice: *Tsc1*^Ctrl^- SD n=5 and *PV-Cre*;*Tsc1^flox/+^*- SD n=7 mice. **S, T**, Examples of trail-by-trail responses evoked by a 10Hz-whisker stimulation vehicle and rapamycin treated control and mutant mice. **U**, **V**, Following 10Hz whisker stimulation, sensory deprived conditional heterozygous and control mice show similar N1 delay reliability (**U**, Mann-Whitney test, p=0.4318) and entrainment (**V**, Welch’s t-test, p=0.8086). Number of mice: Sensory deprived *Tsc1*^Ctrl^- SD n=5 and *PV-Cre*;*Tsc1^flox/+^*- SD n=7 mice. Bar graphs represent mean ± SEM, dots represent individual mouse data. SD: Sensory Deprivation.

Altogether, these data suggest that short-term sensory deprivation during a sensitive postnatal window is sufficient to prevent the development of atypical tactile behavior (exploratory and defensiveness) and aberrant whisker-evoked responses caused by mTORC1 hyperactivity in PV cells.

### Short-term whisker trimming during the third postnatal week prevents the development of sociability deficits in adult *PV-Cre*; *Tsc1^flox/+^* mice

We previously demonstrated that *Tsc1* deletion in PV cells leads to social behavior deficits^17^. Given the essential role of touch in social interaction, atypical sensory processing might underlie these social impairments. Since whisker trimming during the third postnatal week successfully prevented atypical tactile exploratory and avoidance behaviors, we next investigated whether this early-life treatment also impacted social behavior in adulthood. In particular, we tested sociability (preference for a novel mouse vs. a novel object, Figure 9A, B) because this behavior is preserved following whisker deprivation during the third postnatal week. In contrast, social discrimination (preference for a novel vs. familiar mouse) may be impaired under these conditions^42^. We found that, while sham mutant mice did not show a preference for a mouse compared to an object, mutant mice whisker-deprived from P14-21 spent more time investigating the mouse, a behavior comparable to that of control mice (Figure 9 C, D). Therefore, a short sensory experience modulation from P14-21, a period during which mice transition to active whisking^28^, is sufficient to rescue sociability deficits in adult *PVCre;Tsc1^flox/+^* mice.

**Figure 9.**
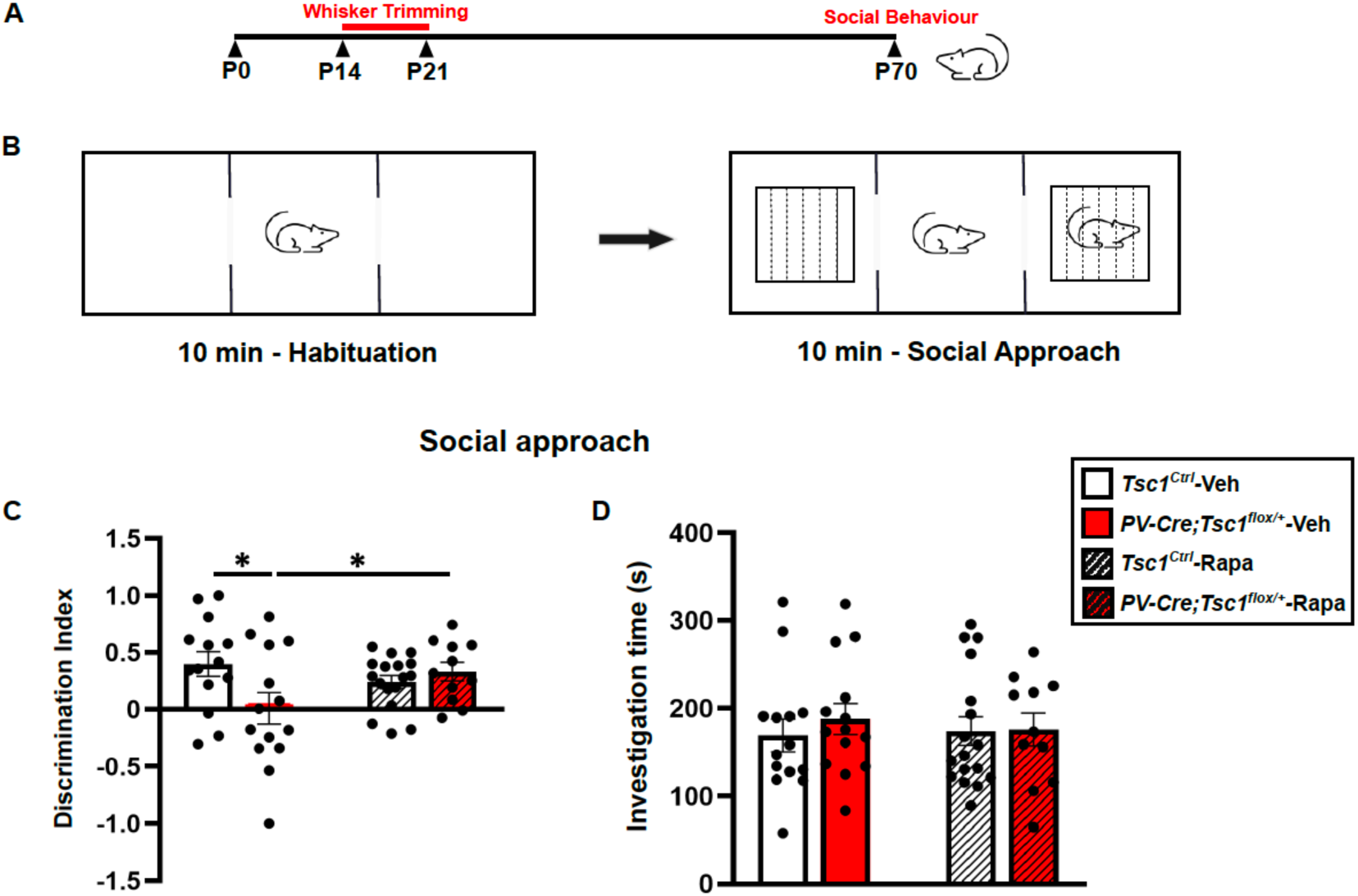
Whisker trimming during the third postnatal week rescues sociability impairment in adult mutant mice. **A**, Schematic of Sensory Deprivation paradigm. **B**, Schematic for testing Social Behavior. **C**, While Sham (NSD) control mice spend more time sniffing a mouse compared to an object, conditional heterozygous sham mice do not show any preference. This phenotype is rescued by whisker deprivation during the third postnatal week (Two-way ANOVA, interaction *p=0.02, Fisher’s LSD multiple comparison test *Tsc1*^Ctrl^**-** NSD vs *PV-Cre;Tsc1^flox/+^***-** NSD, **p=0.0078; *PV-Cre;Tsc1^flox/+^***-** NSD vs *PV-Cre;Tsc1^flox/+^***-** SD, *p=0.0359; *Tsc1*^Ctrl^**-** NSD vs *PV-Cre;Tsc1^flox/+^***-** SD p=0.6585). **D**, No significant differences were noted in total sniffing time amongst all experimental groups (Two-way ANOVA, interaction p=0.6424). Number of non-sensory deprived (NSD) mice: *Tsc1*^Ctrl^ n=14 and *PV-Cre*;*Tsc1^flox/+^* n=14. Number of sensory deprived (SD) mice: *Tsc1*^Ctrl^ n=18 and *PV-Cre*;*Tsc1^flox/+^* n=11. Bar graphs represent mean ± SEM, dots represent individual mouse data.

## DISCUSSION

Up to 90% of individuals with ASD show atypical sensory processing, which is often classified into hyper-responsiveness, hypo-responsiveness, sensory avoidance, and sensory-seeking behaviors^3,43^. Hypofunction of GABAergic interneurons, in particular PV cells, has been suggested to contribute to ASD pathophysiology^9^ and to be causally linked to sensory abnormalities in several mouse models^11–13^. However, it remains unclear whether PV cell dysfunction is sufficient to induce abnormal tactile behaviors or if it represents a form of homeostatic compensation. Here, we investigated how mTORC1 hyper-activation— linked to multiple NDDs with high ASD prevalence— specifically in PV cells affects tactile behavior and whisker-evoked cortical activity, and the underlying cellular mechanisms. Since sensory experience strongly regulates PV circuit maturation during a critical postnatal period^19^, we further explored whether limiting whisker-based touch in the week following the emergence of active exploratory whisking and mature network activity^39^ had an effect on the observed phenotypes.

One of the main discoveries of this study is that mTORC1 overactivation restricted to PV cells is sufficient to drive the emergence of atypical touch-related processing and behaviors. In particular mutant mice showed: 1) increased voluntary exploration of textured objects, 2) enhanced defensiveness during passive whisker stimulation, and 3) altered cortical activity within primary somatosensory cortical circuits, suggestive of increased but less temporally precise touch-related responses. Several mouse models carrying causally-linked risk variants for neurodevelopmental disorders exhibit impaired texture discrimination behaviors^4^. In contrast, to our knowledge here we report for the first time a mouse model showing enhanced voluntary exploration of a different textured object. The findings of the handful of studies that have examined tactile responsivity in ASD are mixed, with some studies reporting higher sensitivity and other reporting hypo-sensitivity^44–46^, which is likely due to wide variations in the type of stimuli used and the site of stimulation. Of note, using a robust psychophysics test, Haigh et al.^47^ reported that individuals with ASD exhibited over-responsiveness to roughness (differences in textures). The behavior we observed in mice with mTORC1 overactivation in PV cells could be due to increased texture discrimination or, rather, tactile-seeking behavior. We favor the interpretation that this behavior reflects tactile seeking rather than enhanced texture discrimination, because mutant mice—unlike their control littermates—maintained an interest in the textured objects for 15–30 minutes following initial exposure, as shown by a higher number of nose-to-object contacts and longer exploration times (Figure 2C, D, Supplemental Figure 3). Tactile seeking has been suggested to be a compensatory mechanism for hyposensitivity^48^. On the other hand, mutant mice also exhibited increased avoidance behaviors when experiencing passive whisker stimulation (Figure 3N), which is consistent with a hyper-sensitivity phenotype. Together, these findings suggest that *Tsc1* haploinsufficiency in PV cells drives context-dependent tactile abnormalities, with active spontaneous exploration and passive imposed stimulation revealing distinct behavioral outcomes. The presence of both behaviors in our mouse model is consistent with data reporting tactile seeking and sensory hypersensitivity as prevalent features in individuals with NDDs/ASD^4^. It is possible that top-down or/and neuromodulatory mechanisms differentially engaged during voluntary exploration versus passive stimulation may shape the specific tactile behavior.

The neurophysiological mechanisms leading to altered tactile behavior are not well understood. In our mouse model, we observed altered temporal processing of tactile stimuli, characterized by delayed N1-ERP latencies, and decreased inter-trial coherence following a 10 Hz whisker stimulation. Consistent with our observations, Miyazaki et al.^49^ reported abnormalities, in particular prolonged peak latencies, in the somatosensory-evoked responses in children with ASD reporting hypersensitivity to tactile stimuli. The reduced temporal precision of repetitive whisker-evoked responses observed in PV cell-specific *Tsc1* haploinsufficient mice is also consistent with reports of heightened behavioral and perceptual variability in ASD, a phenomenon that may rely upon enhanced variability of evoked neural responses^46,47,50–52^. Given that elevated trial-to-trial variability diminishes the reliability of sensory-evoked responses, an alternative hypothesis is that prolonged exploration of the novel-textured object reflects the mouse sensory uncertainty regarding the tactile distinction from the familiar object.

Given that mTORC1 hyperactivation in PV cells impairs their excitatory drive, intrinsic excitability, and output connectivity, we hypothesize that resulting PV cell hypofunction contributes to the observed alterations in whisker-evoked responses. Consistent with this hypothesis, a recent study showed that optogenetically silencing a small fraction of PV cells caused an increase in LFP responses to sensory stimulation, but also a decrease in temporal discrimination of consecutive whisker deflections^53^. Furthermore, in a Fmr1 KO mouse model, chemogenetic PV cell activation significantly increased the number of whisker-responsive, time-locked neurons to levels comparable with wild-type mice^13,41^. Whether increasing PV cell activity by chemogenetic or pharmacological approaches ameliorate the observed phenotypes in adult *PVCre;Tsc1^flox/+^* mice remains to be determined.

The second main finding of this study is that reducing whisker-mediated sensory inputs during the third postnatal week dampens mTORC1 hyperactivation in PV cells, as indicated by pS6 expression levels, and prevents the development of PV cell connectivity deficits, abnormal whisker-evoked cortical activity and atypical tactile behavior caused by *Tsc1* haploinsufficiency in PV cells. In line with our findings, early visual deprivation prevents the hyper-connectivity of PV cells and the decline of visual acuity in a mouse model of Rett’s syndrome^54^. These observations support the hypothesis that abnormal sensory-evoked activity in early life can cause long-term changes in cortical wiring and sensory processing. Notably, apart from reduced PV cell intrinsic excitability (Supplemental figure 8D), we observed no other major phenotypic alterations in whisker-trimmed control mice. It is possible that longer deprivation periods may be required to significantly alter whisker-evoked cortical responses^55^. Alternatively, the effects of whisker-deprivation on control mouse behavior may be more subtle, thus requiring a much larger sample size to be detected.

Our findings are also consistent with a recent study showing that boosting PV cell activity during the third postnatal week (P15-P20), but not earlier (P5-P10), restores network activity within somatosensory cortical circuits and reduces tactile defensiveness behavior in a mouse model of Fragile X syndrome^13^. These data identify a critical developmental window during which PV cell dysfunction has a strong impact on the emergence of atypical tactile behaviors later in life. The end of the second postnatal week coincides with the strengthening of perisomatic GABAergic inhibition onto pyramidal neurons^19,56,57^, but also with the desynchronization of cortical networks^38,39^ and active whisker onset^28^. We hypothesize that, in the presence of mTORC1 hyperactivity, sensory input during this specific maturation phase steers PV cells toward an abnormal developmental path. In accordance with this hypothesis, we found that—similar to whisker-deprivation—treatment with the mTORC1 inhibitor rapamycin during the third postnatal week prevents the development of PV cell connectivity deficits, abnormal whisker-evoked cortical activity and atypical tactile behavior in mutant mice. Notably, during the third postnatal week, PV cell synaptic innervations increase in complexity, via a process that is dependent on neuronal activity and sensory experience^19,37^. Sensory-evoked neuronal activity affects the morphological and functional maturation of PV cells by regulating the expression levels of molecular effectors, including neurotrophic factors^58–60^, extracellular matrix components^61^ and adhesion molecules^62–64^. Several of the signaling pathways recruited by these known molecular regulators of PV cell maturation converges on mTORC1 signaling^33,36^. In *Tsc1* haploinsufficient PV cells, hyper-activated mTORC1 may thus cause sensory signals to be ‘misinterpreted, thus altering the way sensory-evoked activity is integrated during PV cell maturation. This could in turn trigger cellular changes leading to PV circuits hypofunction, ultimately altering touch-related behavior and cortical responses in adulthood. An alternative, though not mutually exclusive, hypothesis is that mTORC1 hyperactivation promotes excessive thalamocortical drive onto PV cells when cortical networks desynchronise. This could trigger excessive feedback inhibition and engage homeostatic mechanisms that ultimately result in PV cell hypoactivity. Consequently, by reducing this excessive drive, whisker deprivation during the third postnatal week may preserve the normal developmental trajectory of these cells.

Regarding the cellular mechanisms leading to PV cell hypofunction, we found that in adult mice, layer 5 PV cells show 1) reduced excitatory drive, consistent with studies showing that mTORC1-mediated signalling regulates glutamatergic synapses formation and plasticity^30^; 2) decreased intrinsic excitability, and 3) reduced perisomatic innervation onto target cells. We focused our analysis on layer 5 PV cells, which are for the most part basket interneurons; however, PV is expressed also by chandelier cells, which target the axon initial segment of pyramidal cells. mTORC1 hyperactivation in those interneurons may contribute to the observed network and behavioral phenotypes. Whether mTORC1 hyperactivation similarly affects the function of chandelier and basket interneurons remain to be determined. Notably, recent studies indicate that mTOR dysregulation does not affect all interneuron subclasses in the same manner. *Tsc1* deletion in SST interneurons was reported to alter molecular identity and promote acquisition of fast-spiking properties rather than reducing firing output^65^. Similarly, deletion of *Tsc1* in VIP interneurons resulted in increased interneuron number and enhanced excitability, associated with altered apoptosis and increased excitatory drive^66^. In contrast, studies examining PV cell circuits in Tsc2^+/−^ and Fmr1^−/−^ models revealed impaired homeostatic regulation of intrinsic excitability rather than enhanced firing^18^. Thus, rather than producing a uniform gain- or loss-of-function phenotype across inhibitory circuits, mTORC1 dysregulation likely impacts distinct interneuron subclasses through cell type–specific mechanisms. It is also possible that the deletion versus the haploinsufficiency of mTORC1 regulators may yield distinct effects. In humans, mutations associated with mTORC1 hyperactivation are mostly dominant; consequently, here we focused our studies in haploinsufficient conditional mice. It remains to be determined whether and to what extent the phenotypes in global *Tsc1* haploinsufficient models resemble those caused by *Tsc1* haploinsufficiency specifically in PV cells.

The third major finding of this study is that whisker trimming during the third postnatal week prevent the development of sociability deficits in adult mice. Since much of our cognitive and social representations are built upon sensory inputs, sensory stimuli and social behaviours have been proposed to reciprocally influence each other throughout development. Supporting this hypothesis, early abnormal sensory sensitivity to stimuli predicts later joint attention and language development^67,68^ and higher levels of social impairment in adults with ASD^69^. Later studies further show that sensory symptoms are predictive of the subsequent appearance of impaired social behaviour and other autistic traits in ASD patients^70^. Touch in particular plays a critical role in several social aspects such as communication, developing social bonds and in brain connectivity^71,72^. Altogether, these studies suggest that abnormal tactile perception may be associated with and contribute to social dysfunctions. Here, we showed that limiting whisker-mediated sensory experience during the third postnatal week was sufficient to prevent sociability deficits in adult mutant mice. This sensitive developmental time window corresponds to the start of exploratory locomotion, which is favored by the maturation of multiple sensory modalities, including eye opening, and the onset of active rhythmic whisking behavior. Our data are consistent with the hypothesis that, in mice with PV cell-restricted *Tsc1* haploinsufficiency, altered cortical circuit activity within the somatosensory cortex disrupt perceptual learning, therefore affecting the development of higher cognitive functions, such as social preference.

Altogether, our findings demonstrate that 1) PV cell impairment leads to atypical tactile processing and behavior, and 2) PV cell maturation involves a sensitive developmental window during which transient interventions can exert enduring effects on both cortical circuitry and behavioral phenotypes. These findings contribute to the growing body of evidence suggesting that specific alterations in PV circuits are linked to convergent pathophysiological phenotypes. Whether, and to what extent, these observations translate to the human condition remains a critical question. While the prolonged postnatal maturation of PV cell connectivity and physiology appears to be a conserved feature^73^, the specific timing and duration of sensitive periods across distinct cortical areas in non-human primates remain largely unexplored. Nonetheless, elucidating how the genetic landscape and environmental factors interact to dictate PV cell maturation and function could provide a framework for novel therapeutic strategies

## ACKNOWLEDGEMENTS

We would like to thank Dr Roberto Araya for his invaluable insights on the manuscript, Kristian Agbogba and Mikael Valton Charette for their technical assistance, Marisol Lavertu-Jolin for helping with automatic quantification of putative synapses, the Comité Institutionnel de Bonne Pratiques Animales en Recherche (CIBPAR), all the personnel of the animal facility of the Research Center of CHU Sainte-Justine (Université de Montreal), Compute Canada and the Plateforme Imagerie Microscopique of the Research Center of CHU Sainte-Justine for their instrumental technical support. We are grateful to all members of our lab for insightful data discussion. We would also like to thank Dr Jozsef Csicsvari for sharing their spike sorting analysis pipeline, Dr Reza Sharif Naeini and Shajenth Premachandran for teaching us the brush test. This work was supported by the Canadian Institutes of Health Research (G.DC), Natural Sciences and Engineering Research Council of Canada (NSERC), La Fondation des Étoiles (G.DC.) and Fonds UdeM pour le partenariat CHU Sainte-Justine -Institut Imagine en épilepsie de l’enfant (G.DC).. C.A.A. is supported by an NSERC fellowship. M.I.C-M and A.C.C are supported by Transforming Autism Care Consortium. R.F. and J.L.D.K. are supported by Fond de Recherche Québec-Santé (FRQS). M.I.C-M and R.F. are supported by The Savoy Foundation.

## REFERENCES

1. Ahn, R.R., Miller, L.J., Milberger, S., McIntosh, D.N. (2004). Prevalence of parents’ perceptions of sensory processing disorders among kindergarten children. Am J Occup Ther 58(3):287–93.

2. Ben-Sasson, A., Carter, A.S., Briggs-Gowan, M.J. (2009). Sensory over-responsivity in elementary school: prevalence and social-emotional correlates. J Abnorm Child Psychol 37(5):705–16.

3. Robertson, C. E., Baron-Cohen S. (2017). Sensory perception in autism. Nat Rev Neurosci 18(11): 671–684.

4. Balasco, L., Provenzano, G., Bozzi, Y. (2019). Sensory Abnormalities in Autism Spectrum Disorders: A Focus on the Tactile Domain, From Genetic Mouse Models to the Clinic. Front Psychiatry 10: 1016.

5. Wiggins, L. D., Robins, D. L., Bakeman, R., Adamson, L. B. (2009). Brief report: sensory abnormalities as distinguishing symptoms of autism spectrum disorders in young children. J Autism Dev Disord 39(7): 1087–1091.

6. Mammen, M. A., Moore, G. A., Scaramella, L. V., Reiss, D., Ganiban, J. M., Shaw, D. S., Leve, L. D., Neiderhiser, J. M. (2015). Infant Avoidance during a Tactile Task Predicts Autism Spectrum Behaviors in Toddlerhood. Infant Ment Health J 36(6): 575–587.

7. LeBlanc, J.J., Fagiolini, M. (2011). "Autism: a "critical period" disorder?". Neural Plast 2011:921680. doi: 10.1155/2011/921680.

8. Iannone, A. F., De Marco Garcia, N. V. (2021). The Emergence of Network Activity Patterns in the Somatosensory Cortex - An Early Window to Autism Spectrum Disorders. Neuroscience 466: 298–309.

9. Contractor, A., I. Ethell. M., Portera-Cailliau C. (2021). Cortical interneurons in autism. Nat Neurosci 24(12): 1648–1659.

10. Sohal, V. S., Rubenstein, J. L. R. (2019). Excitation-inhibition balance as a framework for investigating mechanisms in neuropsychiatric disorders. Mol Psychiatry 24(9): 1248–1257.

11. Goel, A., Cantu, D. A., Guilfoyle, J., Chaudhari, G. R., Newadkar, A., Todisco, B., de Alba, D., Kourdougli, N., Schmitt, L. M., Pedapati, E., Erickson, C. A., Portera-Cailliau, C. (2018). Impaired perceptual learning in a mouse model of Fragile X syndrome is mediated by parvalbumin neuron dysfunction and is reversible. Nat Neurosci 21(10): 1404–1411.

12. Chen, Q., Deister, C. A., Gao, X., Guo, B., Lynn-Jones, T., Chen, N., Wells, M. F., Liu, R., Goard, M. J., Dimidschstein, J., Feng, S., Shi, Y., Liao, W., Lu, Z., Fishell, G., Moore, C. I., Feng G. (2020). Dysfunction of cortical GABAergic neurons leads to sensory hyper-reactivity in a Shank3 mouse model of ASD. Nat Neurosci 23(4): 520–532.

13. Kourdougli, N., Suresh, A., Liu, B., Juarez, P., Lin, A., Chung, D. T., Graven Sams, A., Gandal, M. J., Martinez-Cerdeno, V., Buonomano, D. V., Hall, B. J., Mombereau, C., Portera-Cailliau, C. (2023). Improvement of sensory deficits in fragile X mice by increasing cortical interneuron activity after the critical period. Neuron 111(18): 2863–2880.

14. Karalis, V., Bateup, H. S. (2021). Current Approaches and Future Directions for the Treatment of mTORopathies. Dev Neurosci 1–16.

15. Winden, K.D., Ebrahimi-Fakhari, D., Sahin, M. (2018). Abnormal mTOR Activation in Autism. Annu Rev Neurosci. 41:1–23.

16. Curatolo, P., Moavero, R., de Vries P.J. (2015). Neurological and neuropsychiatric aspects of tuberous sclerosis complex. Lancet Neurol. 14(7):733–45.

17. Amegandjin, C. A., Choudhury, M., Jadhav, V., Carrico, J. N., Quintal, A., Berryer, M., Snapyan, M., Chattopadhyaya, B., Saghatelyan, A., Di Cristo, G. (2021). Sensitive period for rescuing parvalbumin interneurons connectivity and social behavior deficits caused by TSC1 loss. Nat Commun 12(1): 3653.

18. Monday, H.R., Nieto, A.M., Yohannes, S.A., Luxu, S., Wong, K.W., Bolio, F.E., Feldman, D.E. (2025). Physiological and molecular impairment of PV circuit homeostasis in mouse models of autism. bioRxiv [Preprint]. doi: 10.1101/2025.01.08.632056.

19. Chattopadhyaya, B., Di Cristo, H., Higashiyama, H., Knott, G. W., Kuhlman, S. J., Welker, E., Huang, Z. J. (2004). Experience and activity-dependent maturation of perisomatic GABAergic innervation in primary visual cortex during a postnatal critical period. J Neurosci 24(43): 9598–9611.

20. Katagiri, H., Fagiolini, M., Hensch, T.K. (2007). Optimization of somatic inhibition at critical period onset in mouse visual cortex. Neuron 53(6):805–12.

21. Jiao, Y., Zhang, Z., Zhang, C., Wang, X., Sakata, K., Lu, B., Sun, QQ (2011). A key mechanism underlying sensory experience-dependent maturation of neocortical GABAergic circuits in vivo. Proc Natl Acad Sci U S A. 108(29):12131–6.

22. Wu, H. P., Ioffe, J. C., Iverson, M. M., Boon, J. M., Dyck, R. H. (2013). Novel, whisker-dependent texture discrimination task for mice. Behav Brain Res 237: 238–242.

23. Orefice, L. L., Zimmerman, A. L., Chirila, A. M., Sleboda, S. J., Head, J. P., Ginty, D. D. (2016). Peripheral Mechanosensory Neuron Dysfunction Underlies Tactile and Behavioral Deficits in Mouse Models of ASDs. Cell 166(2): 299–313.

24. He, Y., Li, J., Zheng, W., Liu, J., Dong, Z., Yang, L., Tang, S., Zou, Y., Gao, T., Yang, Y., Mo, Z., Wang, S., He, Y., Tang, C., Luo, J., Zhao, J., Guo, G., Li, H., Xiao, L. (2025). Hypomyelination in autism-associated neuroligin-3 mutant mice impairs parvalbumin interneuron excitability, gamma oscillations, and sensory discrimination. Nat Commun. 16(1):6382.

25. Beaulieu-Laroche, L., et al. (2020). TACAN Is an Ion Channel Involved in Sensing Mechanical Pain. Cell 180(5): 956–967 e917.

26. Cheng, L., Duan, B.. Huang, T., Zhang, Y., Chen, Y., Britz, O., Garcia-Campmany, L., Ren, X., Vong, L., Lowell, B. B., Goulding, M., Wang, Y., Ma, Q. (2017). Identification of spinal circuits involved in touch-evoked dynamic mechanical pain. Nat Neurosci 20(6): 804–814.

27. Xu, Q., Tam, M., Anderson, S. A. (2008). Fate mapping Nkx2.1-lineage cells in the mouse telencephalon. J Comp Neurol 506(1): 16–29.

28. Arakawa, H., Erzurumlu, R. S. (2015). Role of whiskers in sensorimotor development of C57BL/6 mice. Behav Brain Res 287: 146–155.

29. Tavazoie, S. F., Alvarez, V. A., Ridenour, D. A., Kwiatkowski, D. J., Sabatini, B. L. (2005). Regulation of neuronal morphology and function by the tumor suppressors Tsc1 and Tsc2. Nat Neurosci 8(12): 1727–1734.

30. Bateup, H. S., Takasaki, K. T., Saulnier, J. L., Denefrio, C. L., Sabatini, B. L. (2011). Loss of Tsc1 in vivo impairs hippocampal mGluR-LTD and increases excitatory synaptic function. J Neurosci 31(24): 8862–8869.

31. Clemens, A.M., Lenschow, C., Beed, P., Li, L., Sammons, R., Naumann, R.K., Wang, H., Schmitz, D., Brecht, M. (2019). Estrus-Cycle Regulation of Cortical Inhibition. Curr Biol. 29(4):605–615.e6.

32. Clayton, K.K., Awwad, B., McGill, M., Stecyk, K.S., Kremer, C., Sudana, K., Skerleva, D., Narayanan, D.P., Zhu, J., Hancock, K.E., Kujawa, S.G., Kozin, E.D., Polley, D.B. (2026). Cortical PV interneurons regulate loudness perception and sustainably reverse loudness hypersensitivity. Neuron. 114(2):325–342.e7.

33. Bernard, C., Exposito-Alonso, D., Selten, M., Sanalidou, S., Hanusz-Godoy, A., Aguilera, A., Hamid, F., Oozeer, F., Maeso, P., Allison, L., Russell, M., Fleck, R. A., Rico, B., Marin, O. (2022). Cortical wiring by synapse type-specific control of local protein synthesis. Science 378(6622): eabm7466.

34. Hou, L., Klann, E. (2004). Activation of the phosphoinositide 3-kinase-Akt-mammalian target of rapamycin signaling pathway is required for metabotropic glutamate receptor-dependent long-term depression. J. Neurosci. 24:6352–61.

35. Tsokas, P., Grace, E.A., Chan, P., Ma, T., Sealfon, S.C., Iyengar, R., Landau, E.M., Blitzer, R.D. (2005). Local protein synthesis mediates a rapid increase in dendritic elongation factor 1A after induction of late long-term potentiation. J Neurosci. 25(24):5833–43.

36. Schratt, G.M., Nigh, E.A., Chen, W.G., Hu, L., Greenberg, M.E. (2004). BDNF regulates the translation of a select group of mRNAs by a mammalian target of rapamycin-phosphatidylinositol 3-kinase-dependent pathway during neuronal development. J Neurosci. 24(33):7366–77.

37. Baho, E., Di Cristo, G. (2012). Neural activity and neurotransmission regulate the maturation of the innervation field of cortical GABAergic interneurons in an age-dependent manner. J Neurosci 32(3):911–8.

38. Ferrer, C., De Marco García, N.V. (2022). The Role of Inhibitory Interneurons in Circuit Assembly and Refinement Across Sensory Cortices. Front Neural Circuits 16:866999.

39. Wu, M.W., Kourdougli, N, Portera-Cailliau, C. (2024). Network state transitions during cortical development. Nat Rev Neurosci. 25(8):535–552.

40. Tsai, P.T., et al (2012). Autistic-like behaviour and cerebellar dysfunction in Purkinje cell Tsc1 mutant mice. Nature. 488:647–651.

41. He, C.X., Cantu, D.A., Mantri, S.S., Zeiger, W.A., Goel, A., Portera-Cailliau, C. (2017). Tactile Defensiveness and Impaired Adaptation of Neuronal Activity in the *Fmr1* Knock-Out Mouse Model of Autism. J Neurosci 37(27):6475–6487.

42. Pan, L., Zheng, L., Wu, X. et al.(2022). A short period of early life oxytocin treatment rescues social behavior dysfunction via suppression of hippocampal hyperactivity in male mice. Mol Psychiatry 27: 4157–4171.

43. Rogers, S. J., Hepburn, S., Wehner, E. (2003). Parent reports of sensory symptoms in toddlers with autism and those with other developmental disorders. J Autism Dev Disord 33(6): 631–642.

44. Blakemore, S.J., Tavassoli, T., Calò, S., Thomas, R.M., Catmur, C., Frith, U., Haggard, P (2006). Tactile sensitivity in Asperger syndrome. Brain Cogn 61(1):5–13.

45. Tommerdahl, M., Tannan, V., Cascio, C.J., Baranek, G.T., Whitsel, B.L (2007). Vibrotactile adaptation fails to enhance spatial localization in adults with autism. Brain Res 1154:116–23.

46. Puts, N. A., Wodka, E. L., Tommerdahl, M., Mostofsky, S. H., Edden, R. A. (2014). Impaired tactile processing in children with autism spectrum disorder. J Neurophysiol 111(9): 1803–1811.

47. Haigh, S. M., Minshew, N., Heeger, D. J., Dinstein, I., Behrmann, M. (2016). Over-Responsiveness and Greater Variability in Roughness Perception in Autism. Autism Res 9(3): 393–402.

48. Piccardi, E.S., Begum, Ali. J., Jones, E.J.H., Mason, L., Charman, T., Johnson, M.H., Gliga. T.; BASIS/STAARS Team (2021). Behavioural and neural markers of tactile sensory processing in infants at elevated likelihood of autism spectrum disorder and/or attention deficit hyperactivity disorder. J Neurodev Disord 13(1):1.

49. Miyazaki, M., Fujii, E., Saijo, T., Mori, K., Hashimoto, T., Kagami, S., Kuroda, Y. (2007). Short-latency somatosensory evoked potentials in infantile autism: evidence of hyperactivity in the right primary somatosensory area. Dev Med Child Neurol 49(1): 13–17.

50. Milne, E. (2011). Increased intra-participant variability in children with autistic spectrum disorders: evidence from single-trial analysis of evoked EEG. Front Psychol 2: 51.

51. Dinstein, I., Heeger, D. J., Lorenzi, L., Minshew, N. J., Malach, R., Behrmann, M. (2012). Unreliable evoked responses in autism. Neuron 75(6): 981–991.

52. Haigh, S. M., Heeger, D. J., Dinstein, I., Minshew, N., Behrmann, M. (2015). Cortical variability in the sensory-evoked response in autism. J Autism Dev Disord 45(5): 1176–1190.

53. Yeganeh, F., Knauer, B., Guimaraes Backhaus, R., Yang, J. W., Stroh, A., Luhmann, H. J., Stuttgen, M. C. (2022). Effects of optogenetic inhibition of a small fraction of parvalbumin-positive interneurons on the representation of sensory stimuli in mouse barrel cortex. Sci Rep 12(1): 19419.

54. Durand, S., Patrizi, A., Quast, K. B., Hachigian, L., R. Pavlyuk, R., Saxena, A., Carninci, P., Hensch, T. K., Fagiolini, M. (2012). NMDA receptor regulation prevents regression of visual cortical function in the absence of Mecp2. Neuron 76(6): 1078–1090.

55. Shoykhet, M., Land, P.W., Simons, D.J (2005). Whisker trimming begun at birth or on postnatal day 12 affects excitatory and inhibitory receptive fields of layer IV barrel neurons. J Neurophysiol 94(6):3987–95.

56. Favuzzi, E., Deogracias, R., et al (2019). Distinct molecular programs regulate synapse specificity in cortical inhibitory circuits. Science 363(6425):413–417.

57. Gour, A., Boergens, K.M., Heike, N., Hua, Y., Laserstein, P., Song, K., Helmstaedter, M. (2021). Postnatal connectomic development of inhibition in mouse barrel cortex. Science 371(6528):eabb4534.

58. Huang, Z.J., Kirkwood, A., Pizzorusso, T., Porciatti, V., Morales, B., Bear, M.F., Maffei, L., Tonegawa, S. (1999). BDNF regulates the maturation of inhibition and the critical period of plasticity in mouse visual cortex. Cell 98(6):739–55.

59. Baho, E., Chattopadhyaya, B., Lavertu-Jolin, M., Mazziotti, R., Awad, P.N., Chehrazi, P., Groleau, M., Jahannault-Talignani, C., Vaucher, E., Ango, F., Pizzorusso, T., Baroncelli, L., Di Cristo, G. (2019). p75 Neurotrophin Receptor Activation Regulates the Timing of the Maturation of Cortical Parvalbumin Interneuron Connectivity and Promotes Juvenile-like Plasticity in Adult Visual Cortex. J Neurosci 39(23):4489–4510.

60. Chehrazi, P., Lee, K. K. Y., Lavertu-Jolin, M., Abbasnejad, Z., Carreno-Munoz, M. I., Chattopadhyaya, B., Di Cristo, G. (2023). The p75 Neurotrophin Receptor in Preadolescent Prefrontal Parvalbumin Interneurons Promotes Cognitive Flexibility in Adult Mice. Biol Psychiatry 94(4): 310–321.

61. Favuzzi, E., Marques-Smith, A., Deogracias, R., Winterflood, C.M., Sánchez-Aguilera, A., Mantoan, L., Maeso, P., Fernandes, C., Ewers, H., Rico, B. (2017). Activity-Dependent Gating of Parvalbumin Interneuron Function by the Perineuronal Net Protein Brevican. Neuron 95(3):639–655.

62. Di Cristo, G., Chattopadhyaya, B., Kuhlman, S.J., Fu, Y., Bélanger, M.C., Wu, C.Z., Rutishauser, U., Maffei, L., Huang, Z.J. (2007). Activity-dependent PSA expression regulates inhibitory maturation and onset of critical period plasticity. Nat Neurosci 10(12):1569–77.

63. Del Pino, I., García-Frigola, C., Dehorter. N., et al (2013). Erbb4 deletion from fast-spiking interneurons causes schizophrenia-like phenotypes. Neuron 79(6):1152–68.

64. Chattopadhyaya, B., Baho, E., Huang, Z.J., Schachner, M., Di Cristo, G. (2013). Neural cell adhesion molecule-mediated Fyn activation promotes GABAergic synapse maturation in postnatal mouse cortex. J Neurosci 33(14):5957–68.

65. Malik, R., Pai, E.L., Rubin, A.N., Stafford, A.M., Angara, K., Minasi, P., Rubenstein, J.L., Sohal, V.S., Vogt, D. (2019). Tsc1 represses parvalbumin expression and fast-spiking properties in somatostatin lineage cortical interneurons. Nat Commun 10(1):4994.

66. Hu, J.S., Malik, R., Sohal, V.S., Rubenstein, J.L., Vogt, D. (2023). *Tsc1* Loss in VIP-Lineage Cortical Interneurons Results in More VIP+ Interneurons and Enhanced Excitability. Cells 13(1):52.

67. Boyd, B. A., Woodard, C. R., Bodfish, J. W. (2013). Feasibility of exposure response prevention to treat repetitive behaviors of children with autism and an intellectual disability: a brief report. Autism 17(2): 196–204.

68. Estes, A., Zwaigenbaum, L., et al. (2015). Behavioral, cognitive, and adaptive development in infants with autism spectrum disorder in the first 2 years of life. J Neurodev Disord 7(1): 24.

69. Hilton, C. L., Harper, J. D., Kueker, R. H., Lang, A. R., Abbacchi, A. M., Todorov, A., LaVesser, P. D. (2010). Sensory responsiveness as a predictor of social severity in children with high functioning autism spectrum disorders. J Autism Dev Disord 40(8): 937–945.

70. Mammen, M. A., Moore, G. A., Scaramella, L. V., Reiss, D., Ganiban, J. M., Shaw, D. S., Leve, L. D., Neiderhiser, J. M. (2015). Infant Avoidance during a Tactile Task Predicts Autism Spectrum Behaviors in Toddlerhood. Infant Ment Health J 36(6): 575–587.

71. Dunbar, R. I. (2010). The social role of touch in humans and primates: behavioural function and neurobiological mechanisms. Neurosci Biobehav Rev 34(2): 260–268.

72. Brauer, J., Xiao, Y., Poulain, T., Friederici, A. D., Schirmer, A. (2016). Frequency of Maternal Touch Predicts Resting Activity and Connectivity of the Developing Social Brain. Cereb Cortex 26(8): 3544–3552.

73. Hosseini Fin, N.S., Yip, A., Scott, J.T., Teo, L., Homman-Ludiye, J., Bourne, J.A. (2025). Developmental dynamics of marmoset prefrontal cortical SST and PV interneuron networks highlight primate-specific features. Development 152(10):dev204254.

74. Sousa, V. H., Miyoshi, G., Hjerling-Leffler, J., Karayannis, T., Fishell, G. (2009). Characterization of Nkx6-2-derived neocortical interneuron lineages. Cereb Cortex 19 **Suppl 1**(Suppl 1): i1–10.

75. Tremblay, M. E., Riad, M., Bouvier, D., Murai, K. K., Pasquale, E. B., Descarries, L., Doucet, G. (2007). Localization of EphA4 in axon terminals and dendritic spines of adult rat hippocampus. J Comp Neurol 501(5): 691–702.

76. Carreno-Munoz, M. I., Chattopadhyaya, B., Agbogba, K., Cote, V., Wang, S., Levesque, M., Avoli, M., Michaud, J. L., Lippé, S., Di Cristo G. (2022). Sensory processing dysregulations as reliable translational biomarkers in SYNGAP1 haploinsufficiency. Brain 145(2): 754–769.

77. Csicsvari, J., Hirase, H., Czurko, A., Buzsaki, G. (1998). Reliability and state dependence of pyramidal cell-interneuron synapses in the hippocampus: an ensemble approach in the behaving rat. Neuron 21(1): 179–189.

78. Batista-Brito, R., Majumdar, A., Nuno, A., Ward, C., Barnes, C., Nikouei, K., Vinck, M., Cardin, J. A. (2023). Developmental loss of ErbB4 in PV interneurons disrupts state-dependent cortical circuit dynamics. Mol Psychiatry 28(7):3133–3143.

