## Supplemental Data for "Sensory experience and mTORC1 interplay orchestrates the maturation of cortical interneuron connectivity and tactile sensitivity"

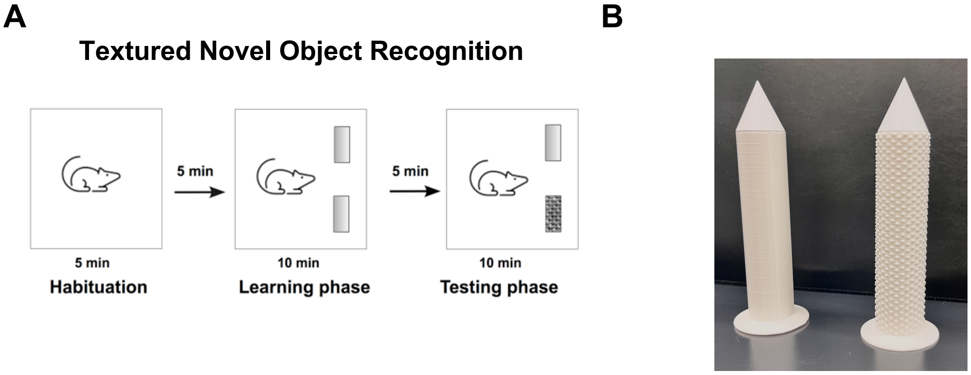


**Supplemental Figure 1. Textured novel object discrimination test. A,** Schematic representation of texture novel object recognition test. **B**, Objects used for the texture novel object recognition test.

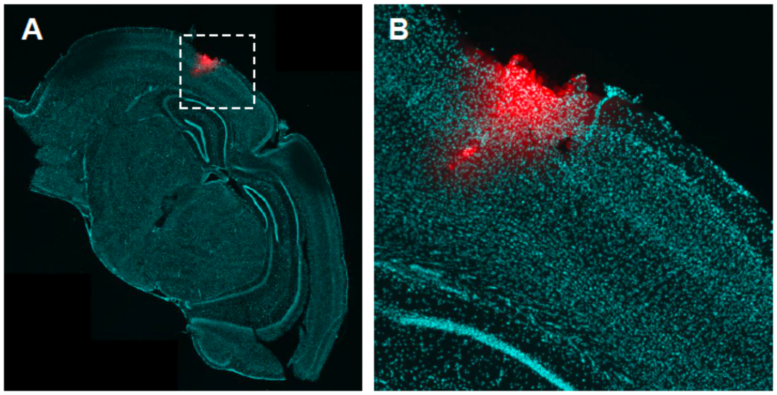


**Supplemental Figure 2. LFP recording approach. A, B,** Low (**A**) and high (**B**) magnification images showing electrode positioning. The electrode was dipped in DiI (red). Cell nuclei are labeled with DAPI (blue).

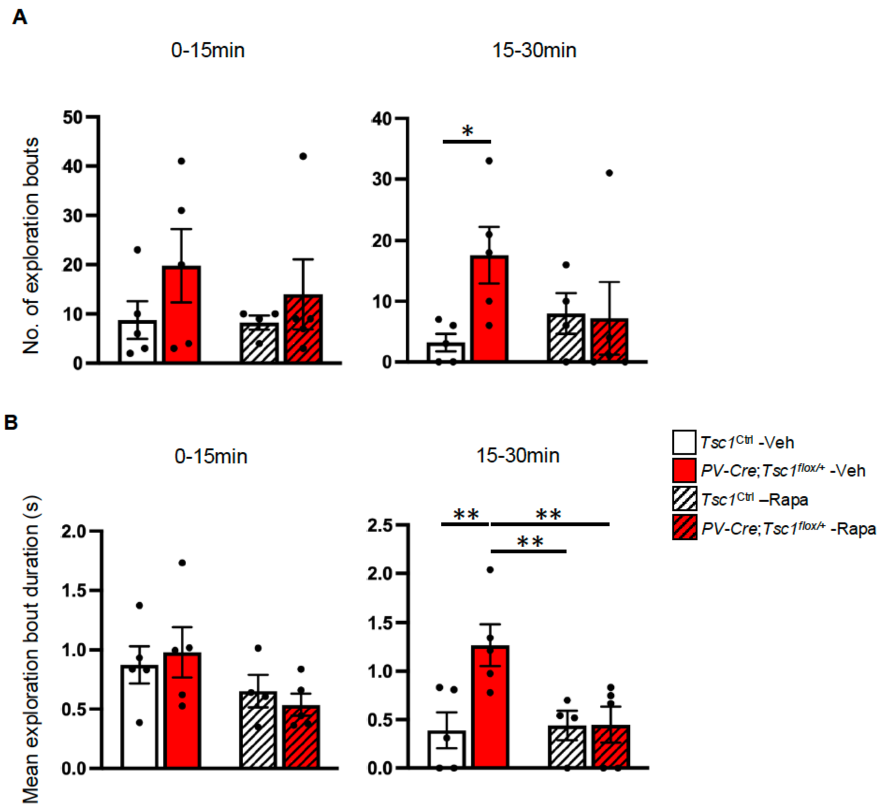


**Supplemental Figure 3. Mutant mice show increased number and mean duration of exploration bouts of textured objects compared to control littermates. A, B,** Number of exploration bouts (**A**, nose-to-object contacts) and duration of each exploration bout (**B**) for vehicle and rapamycin-treated control and mutant mice. While mutant and control mice exhibited comparable exploration behavior during the first 15 minutes (**A**, 0-15': Two-way ANOVA, Fisher LSD test, interaction p=0.66; **B**, 0-15': Two-way ANOVA, Fisher LSD test, interaction p=0.496); mutant mice showed significantly higher number of exploration bouts, along with increased single-bout exploration time, from 15–30 minutes (**A**, 15-30': Two-way ANOVA, Fisher LSD test, interaction p=0.10, *Tsc1*$^{\mathrm{Ctrl}}$-Veh vs *PV-Cre*;*Tsc1*$^{flox/+}$-Veh *p=0.0292; **B**, 15-30': Two-way ANOVA, Fisher LSD test, interaction p=0.0378, *Tsc1*$^{\mathrm{Ctrl}}$-Veh vs *PV-Cre*;*Tsc1*$^{flox/+}$-Veh **p=0.0043, *PV-Cre*;*Tsc1*$^{flox/+}$-Veh vs *PV-Cre*;*Tsc1*$^{flox/+}$-Rapa **p=0.0069, *PV-Cre*;*Tsc1*$^{flox/+}$-Veh vs *Tsc1*$^{\text{Ctrl}}$-Rapa **p=0.0092). Number of mice: Vehicle-treated *Tsc1*$^{\text{Ctrl}}$ n=5 and *PV-Cre*;*Tsc1*$^{flox/+}$ n=5; Rapamycin-treated *Tsc1*$^{\text{Ctrl}}$ n=4 and *PV-Cre*;*Tsc1*$^{flox/+}$ n=5. Bar graphs represent mean ± SEM, dots represent individual mouse data.

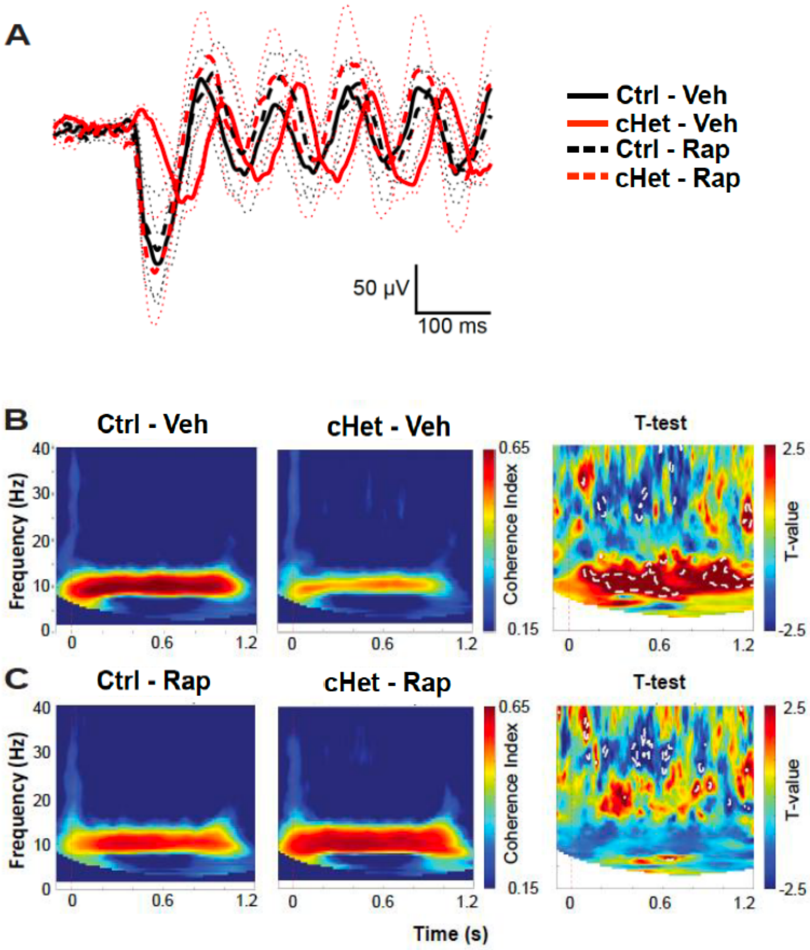


**Supplemental Figure 4. Rapamycin treatment during the third postnatal week rescues evoked ERP dynamics and entrainment in awake adult mutant mice. A**, Superposed grand average traces of ERPs evoked by 10 Hz whisker-deflection and recorded in layer 5 of somatosensory cortex. Note that in vehicle-treated cHet mice the ERP delay is longer than in the other experimental groups. **B, C,** Inter-trial coherence and Welch's t-test time-frequency maps showing comparison between vehicle-treated (**B**) and rapamycin-treated (**C**) control and conditional heterozygous mice. Statistical differences (P<0.05) are marked by white dotted lines. Number of mice: Vehicle-treated *Tsc1*$^{\mathrm{Ctrl}}$ n=6 and *PV-Cre*;*Tsc1*$^{flox/+}$ n=5; Rapamycin-treated *Tsc1*$^{\mathrm{Ctrl}}$ n=5 and *PV-Cre*;*Tsc1*$^{flox/+}$ n=5.

**A**

| | PV cells in Tsc1$^{Ctrl}$ (n=23) 5 animals | PV cells in *PV-Cre;Tsc1*$^{flox/+}$ (n=21) 5 animals | LMM |
|---|---|---|---|
| Resting membrane potential (mV) | -67.5 ± 1.2 | -66.1 ± 1.0 | F=0.769 p=0.385 |
| Input resistance (MΩ ) | 160.2 ± 11.7 | 204.2 ± 20.9 | F=0.455 p=0.504 |
| Membrane capacitance (pF) | 45-0 ± 6.8 | 63.1 ± 10.0 | F=1.823 p=0.184 |
| Membrane time constant (ms) | 13.6 ± 0.9 | 15.8 ± 1.6 | F=0.564 p=0.457 |
| Rheobase current (pA) | 193.0 ± 14.4 | 150.5 ± 16.8 | F=1.072 p=0.306 |
| AP threshold (mV) | -45.1 ± 1.5 | -49.3 ± 1.7 | F=0.881 p=0.353 |
| AP amplitude (mV) | 63.0 ± 1.8 | 71.0 ± 2.1 | F=3.097 p=0.086 |
| AP latency (ms) | 41.0 ± 7.9 | 46.4 ± 20.7 | F=0.066 p=0.798 |
| AP half width (ms) | 0.63 ± 0.02 | 0.74 ± 0.03 | F=2.013 p=0.163 |
| fAHP (mV) | -13.3 ± 0.6 | -13.2 ± 0.7 | F=0.005 p=0.942 |
| fAHP (ms) | 2.78 ± 0.12 | 2.93 ± 0.12 | F=0.285 p=0.596 |
| $I_h$ sag (mV) | 1.14 ± 0.11 | 1.28 ± 0.15 | F=0.334 p=0.566 |

**B**

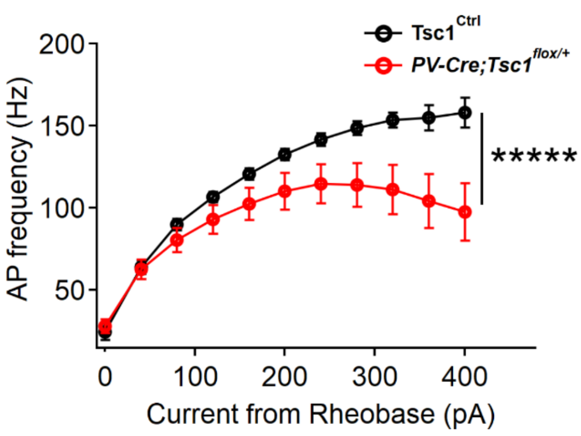


**Supplemental Figure 5. Layer 5 PV cells show reduced firing in adult PV cell-specific *Tsc1* haploinsufficient mice. A,** Summary table describing the intrinsic properties of layer 5 PV cells recorded in somatosensory cortex of adult (P60-85) *Tsc1*$^{Ctrl}$ and *PV-Cre*;*Tsc1*$^{flox/+}$ mice. LMM: linear mixed model **B**, Frequency–current (F–I) relationship showing AP firing frequency in response to 500-ms depolarizing current steps (40 pA increments) plotted relative to rheobase. Layer 5 PV cells from *PV-Cre;Tsc1*$^{flox/+}$ (red line) exhibit reduced firing compared to *Tsc1*$^{Ctrl}$ PV cells (black line) across increasing current injections (Two-way repeated measures ANOVA with Sidak's multiple comparison *post hoc* test, *****p<0.00001). *Tsc1*$^{Ctrl}$: n=23 PV cells from 5 mice; *PV-Cre*;*Tsc1*$^{flox/+}$: n=21 from 5 mice. Lines represent mean ± SEM.

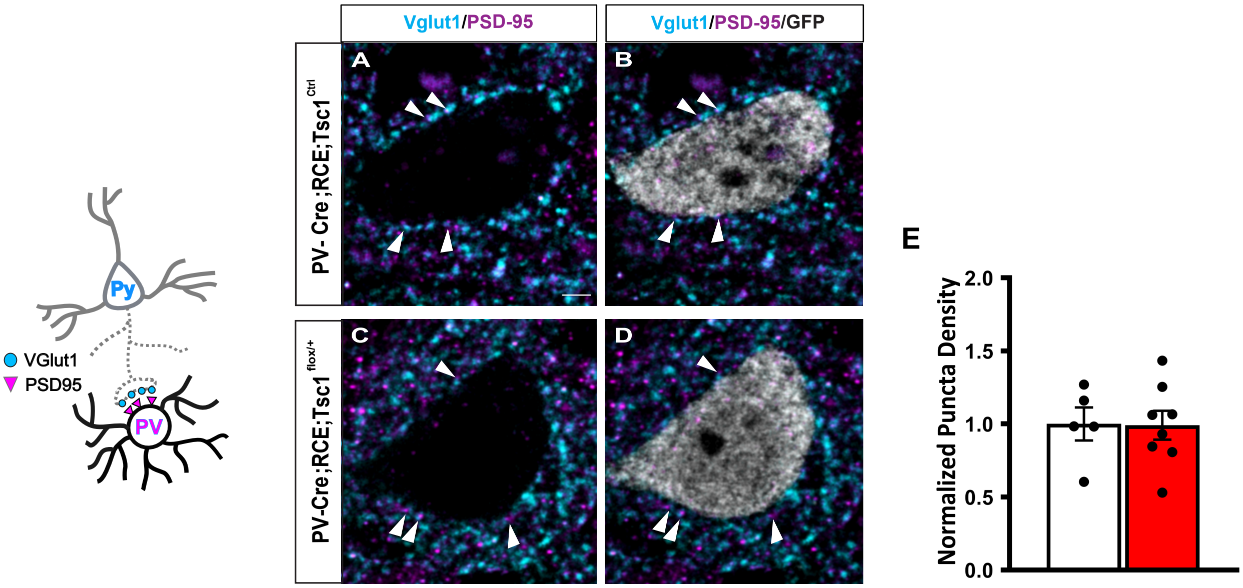


**Supplemental Figure 6. Glutamatergic inputs onto PV cell somata are not altered by *Tsc1* haploinsufficiency in preadolescent mice. A-D,** Representative immunostained sections of somatosensory cortex labeled for Vglut1/PSD-95 (cyan/magenta) and GFP (gray) in P22 *Tsc1*$^{\text{Ctrl}}$ (**A**, **B**) and *PV-Cre;Tsc1*$^{flox/+}$ (**C**, **D**) mice. **E**, Vglut1/PSD-95 colocalized puncta density around layer 5 PV cell somata is not different between the two genotypes (Welch's t-test, p=0.9543). Number of mice: n=5 for *Tsc1*$^{\text{Ctrl}}$ and n=8 for *PV-Cre;RCE;Tsc1*$^{flox/+}$.

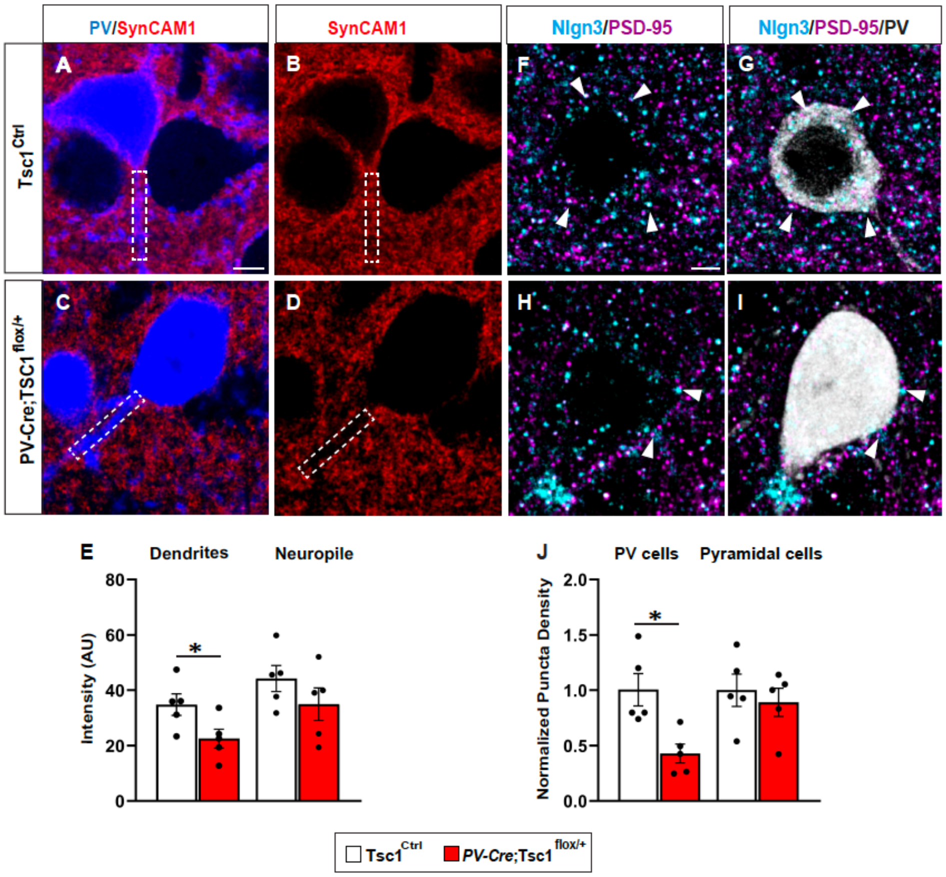


**Supplemental Figure 7. SynCAM1 and Nlg3 expression levels are decreased in *Tsc1* haploinsufficient PV cells in preadolescent mice. A-D**, Representative immunostained sections of somatosensory cortex labeled for SynCAM1 (red), and PV (blue) in ~P25 $Tsc1^{\text{Ctrl}}$ (**A**, **B**) and *PV-Cre;Tsc1*$^{flox/+}$ (**C**, **D**) mice. Examples of PV cell dendrites are outlined by dotted white line. **E**, SynCAM1 expression levels are significantly reduced in PV cell dendrites (Welch's t-test, *p=0.0459), but not in the surrounding neuropile (Welch's t-test, p=0.2564) in layer 5 of the somatosensory cortex of mutant mice compared to control littermates. **F-G**, Representative immunostained sections of somatosensory cortex labeled for Nlgn3 (cyan), PSD-95 (magenta) and PV (gray) in ~P25 $Tsc1^{\text{Ctrl}}$ (**F**, **G**) and *PV-Cre;Tsc1*$^{flox/+}$ (**H**, **I**) mice. White arrowheads indicate Nlgn3/PSD-95 colocalized puncta. **J**, The density of Nlgn3 puncta colocalized with PSD95 onto layer 5 PV cell somata (Welch's t-test, *p=0.0459), but not onto nearby pyramidal neuron somata (Welch's t-test, p=0.5862), is reduced in conditional heterozygous mice. Scale bar: 5 µm. Number of mice: n=5 for both genotypes. Scale bar: 5 µm. Bar graphs represent mean ± SEM. Dots represent single mouse data.

**A**

| | PV cells in *PV-Cre;Tsc1*$^{flox/+}$ - NSD (n=23) 4 animals | PV cells in *PV-Cre;Tsc1*$^{flox/+}$ - SD (n=31) 5 animals | LMM |
|---|---|---|---|
| Resting membrane potential (mV) | -68.6 ± 1.2 | -68.1 ± 0.9 | F=0.092 p=0.763 |
| Input resistance (MΩ ) | 190.4 ± 22.7 | 150.2 ± 11.1 | F=2.371 p=0.130 |
| Membrane capacitance (pF) | 31.7 ± 3.5 | 42.9 ± 6.9 | F=1.565 p=0.217 |
| Membrane time constant (ms) | 11.2 ± 1.0 | 9.1 ± 0.6 | F=3.014 p=0.088 |
| Rheobase current (pA) | 175.6 ± 17.0 | 233.5 ± 21.0 | F=3.339 p=0.073 |
| AP threshold (mV) | -47.3 ± 1.4 | -44.1 ± 0.8 | F=3.429 p=0.070 |
| AP amplitude (mV) | 64.4 ± 2.1 | 61.2 ± 1.5 | F=1.731 p=0.194 |
| AP latency (ms) | 42.4 ± 14.7 | 44.9 ± 14.9 | F=0.014 p=0.905 |
| AP half width (ms) | 0.78 ± 0.05 | 0.60 ± 0.02 | F=4.993 p=0.030* |
| fAHP (mV) | -13.2 ± 0.9 | -14.4 ± 0.6 | F=1.316 p=0.257 |
| fAHP (ms) | 3.27 ± 0.31 | 2.53 ± 0.15 | F=2.459 p=0.123 |
| $I_h$ sag (mV) | 1.14 ± 0.11 | 0.92 ± 0.08 | F=1.634 p=0.207 |

**B**

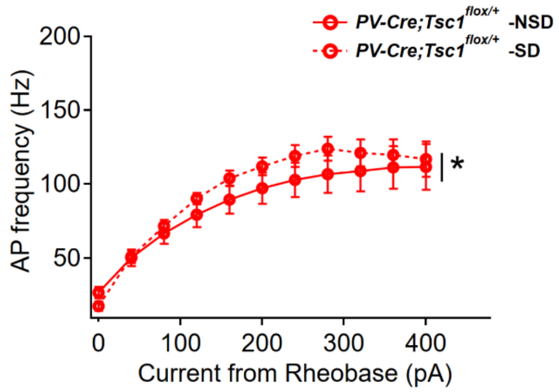


**C**

| | PV cells in Tsc1$^{Ctrl}$-NSD (n=24) 5 animals | PV cells in Tsc1$^{Ctrl}$-SD (n=14) 3 animals | LMM |
|---|---|---|---|
| Resting membrane potential (mV) | -66.8 ± 1.0 | -71.9 ± 1.2 | F=9.396 p=0.004** |
| Input resistance (MΩ ) | 164.8 ± 10.8 | 188.6 ± 16.1 | F=1.697 p=0.201 |
| Membrane capacitance (pF) | 41.8 ± 6.8 | 26.6 ± 1.8 | F=2.768 p=0.105 |
| Membrane time constant (ms) | 13.5 ± 0.9 | 10.8 ± 0.7 | F=2.818 p=0.102 |
| Rheobase current (pA) | 185.0 ± 12.2 | 174.3 ± 13.6 | F=0.331 p=0.569 |
| AP threshold (mV) | -45.5 ± 1.4 | -49.0 ± 1.8 | F=1.391 p=0.246 |
| AP amplitude (mV) | 63.4 ± 1.7 | 61.5 ± 2.7 | F=0.292 p=0.593 |
| AP latency (ms) | 36.2 ± 7.2 | 19.8 ± 1.3 | F=3.084 p=0.088 |
| AP half width (ms) | 0.65 ± 0.02 | 0.70 ± 0.03 | F=1.388 p=0.246 |
| fAHP (mV) | -13.3 ± 0.6 | -14.9 ± 1.2 | F=1.767 p=0.192 |
| fAHP (ms) | 2.88 ± 0.10 | 2.80 ± 0.13 | F=0.235 p=0.631 |
| $I_h$ sag (mV) | 1.19 ± 0.10 | 1.19 ± 0.15 | F=0.006 p=0.949 |

**D**

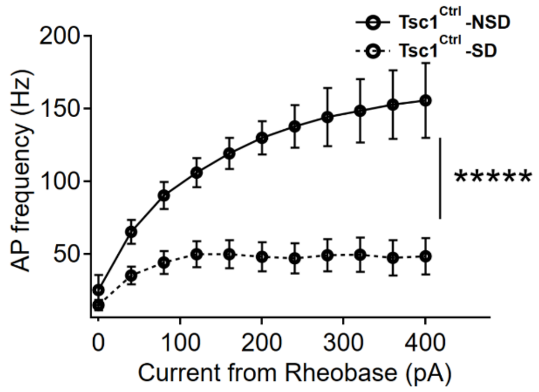


**Supplemental Figure 8. Whisker deprivation during the third postnatal week differentially affects layer 5 PV cells firing in adult control and mutant mice. A, C.** Summary table describing the intrinsic properties of layer 5 PV cells recorded in somatosensory cortex of adult (P60-75) sham (non-sensory deprived, NSD) and whisker deprived (sensory deprived, SD) *Tsc1*$^{Ctrl}$ and *PV-Cre*;*Tsc1*$^{flox/+}$ mice. LMM: linear mixed model. **B, D**. F–I relationship illustrating the effect of whisker deprivation during the third postnatal week on AP evoked firing frequency across increasing depolarizing current injections (500 ms, 40 pA increments), plotted relative to rheobase.

Recordings were obtained from layer 5 PV interneurons of *PV-Cre;Tsc1*$^{flox/+}$ and *Tsc1*$^{\mathrm{Ctrl}}$ mice. **B,** Whisker deprivation significantly increased firing in mutant PV cells from *PV-Cre;Tsc1*$^{flox/+}$-SD (dashed red line) compared to *PV-Cre;Tsc1*$^{flox/+}$-NSD mice (solid red line; two-way repeated measures ANOVA with Sidak's multiple comparison *post hoc* test, *p=0.016; **D,** In contrast, whisker deprivation significantly reduced evoked firing in PV cell from *Tsc1*$^{\mathrm{Ctrl}}$-SD (dashed black line) compared to *Tsc1*$^{\mathrm{Ctrl}}$-NSD mice (solid black line; two-way repeated measures ANOVA with Sidak's multiple comparison *post hoc* test, *****p<0.00001). *PV-Cre*;*Tsc1*$^{flox/+}$-NSD: n=23 PV cells from 4 mice; *PV-Cre*;*Tsc1*$^{flox/+}$-SD: n=31 PV cells from 5 mice; *Tsc1*$^{Ctrl}$-NSD*:* n=24 PV cells from 5 mice; *Tsc1*$^{\mathrm{Ctrl}}$-SD: n=14 PV cells from 3 mice.

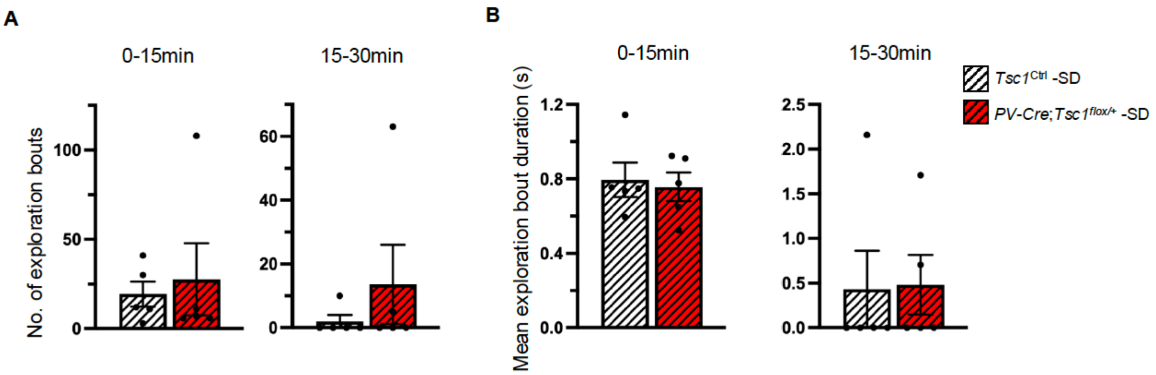


**Supplemental Figure 9. Adult control and mutant mice whisker-trimmed during the third postnatal week show comparable exploration time of textured objects. A, B,** Number of exploration bouts (**A**, nose-to-object contacts) and duration of each exploration bout (**B**). **A**, Mann-Whitney test, 0-15’ p=0.5873. 15-30’ p=0.7222; **B**, Mann-Whitney test, 0-15’ p>0.99. 15-30’ p>0.99. Number of mice: Sensory deprived *Tsc1*$^{\mathrm{Ctrl}}$- SD n=5 and *PV-Cre*;*Tsc1*$^{flox/+}$- SD n=5 mice. Bar graphs represent mean ± SEM, dots represent individual mouse data. SD: Sensory Deprivation.

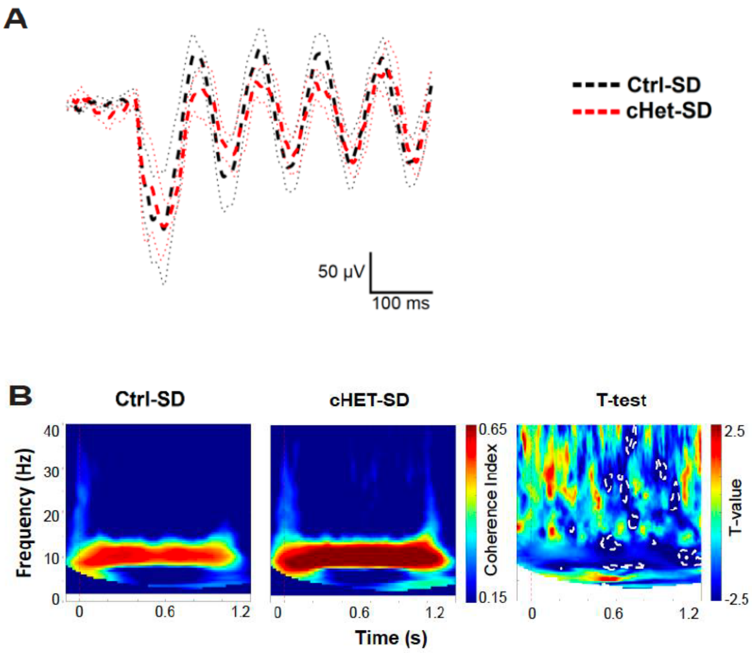


**Supplemental Figure 10. Whisker deprivation during the third postnatal week rescues evoked ERP dynamics and entrainment in adult mutant mice. A**, Superposed grand average traces of ERPs evoked by 10 Hz whisker-deflection and recorded in layer 5 of somatosensory cortex. **B,** Inter-trial coherence and Welch's t-test maps showing comparison between whisker deprived wild-type and conditional heterozygous mice. Statistical differences (P<0.05) are marked by white dotted lines. Number of mice: *Tsc1*$^{\mathrm{Ctrl}}$ -SD n=6 and *PV-Cre*;*Tsc1*$^{flox/+}$ -SD n=5. SD: sensory deprivation.
